# Mechanistic Insights into MYO1C-Mediated Rhodopsin Trafficking and Rod Photoreceptor Homeostasis

**DOI:** 10.64898/2026.09.04.749496

**Authors:** Rakesh Radhakrishnan, Vincent Norton, Heidi Roehrich, Sebahattin Cureoglu, Altaf A. Kondkar, Rafael da Costa Monsanto, Frederik J van Kuijk, Glenn P. Lobo

## Abstract

Rhodopsin trafficking from the photoreceptor inner segment to the outer segment is essential for photoreceptor function, yet the molecular mechanism(s) regulating this process remain incompletely understood. MYO1C is an actin-based motor protein implicated in intracellular cargo trafficking. Here, we investigated its role in rhodopsin trafficking and photoreceptor cell homeostasis. In-silico docking identified a putative interaction between the MYO1C C-terminal region and the C-terminal region of rhodopsin containing the conserved VxPx ciliary trafficking motif. Biochemical studies confirmed that full-length MYO1C interacts with rhodopsin, whereas deletion of the MYO1C C-terminal domain abolished this interaction. Live-cell imaging, ciliary localization, and fluorescence recovery after photobleaching in hTERT-RPE1 cells demonstrated that the MYO1C C-terminal region is required for efficient rhodopsin trafficking, membrane localization, and ciliary targeting. In native murine rod photoreceptors MYO1C localized to both inner and outer segments. Global *Myo1c* deficiency in mice caused age-dependent rhodopsin mislocalization, apo-opsin accumulation and progressive retinal dysfunction, characterized primarily by reduced scotopic ERG responses and delayed a-wave recovery, beginning at 6-months, while photopic responses were relatively preserved. Rod-specific *Myo1c* deletion similarly caused progressive scotopic dysfunction and reduced a-wave recovery following light stimulation. In contrast, cone-specific *Myo1c* deletion preserved photopic function and a-wave recovery. Together, these findings identify MYO1C as an important regulator of rhodopsin trafficking and demonstrate a preferential, cell-autonomous requirement for MYO1C in maintaining rod photoreceptor homeostasis and phototransduction recovery. These findings establish a mechanistic link between MYO1C-dependent rhodopsin trafficking and age-dependent rod photoreceptor cell dysfunction.

## INTRODUCTION

Rhodopsin is the most abundant protein in rod photoreceptor outer segments (OS) and is essential for the initiation of phototransduction and maintenance of photoreceptor structure [1–7]. Newly synthesized rhodopsin is produced in the inner segment (IS) and must be transported through the connecting cilium (CC) to the OS, where it is incorporated into newly forming disc membranes [3,4]. This highly polarized trafficking pathway is essential for photoreceptor cell survival and visual function. Disruption of rhodopsin trafficking results in its accumulation within the IS and photoreceptor cell body and is a characteristic feature of several inherited retinal degenerative diseases, including retinitis pigmentosa (RP) [2,4,6]. Despite the importance of this pathway, the molecular mechanisms that coordinate rhodopsin transport from the IS to the OS, membrane delivery and compartmentalization, and OS targeting remain incompletely understood [2, 4].

Rhodopsin trafficking is mediated by a complex network of molecular motors, adaptor proteins, cytoskeletal elements, and membrane-associated factors [1–3]. The conserved C-terminal VxPx motif of rhodopsin is particularly important for its polarized trafficking from the photoreceptor IS toward the OS [4,6–13]. Interactions between this targeting sequence and components of the intracellular trafficking machinery ensure appropriate delivery of rhodopsin to the photoreceptor OS. Mutations within or surrounding the C-terminal trafficking region of rhodopsin can disrupt its localization and cause retinal degeneration, emphasizing the importance of this pathway for photoreceptor homeostasis [7–13]. However, the molecular motors, cargo-binding proteins, and specific cargo binding protein domain(s), that directly participate in rhodopsin trafficking, movement, and membrane delivery remain incompletely defined.

Myosin proteins are actin-based molecular motors that regulate intracellular cargo transport, membrane organization, and cytoskeletal dynamics [14–17]. MYO1C is a class I myosin that contains an N-terminal ATPase motor domain, a neck region containing IQ1 and IQ2 motifs, and a C-terminal cargo-binding region [17]. MYO1C has established functions in membrane-cytoskeleton interactions and intracellular trafficking, but its role in photoreceptor protein transport has not been well characterized. Given the highly specialized organization of photoreceptors and their dependence on coordinated actin- and membrane-associated trafficking mechanisms, MYO1C represents a potential regulator of rhodopsin transport and membrane delivery [17, 18].

Our previous studies demonstrated that global loss of *Myo1c* in mice results in abnormal rhodopsin localization, with a significant proportion of rhodopsin mislocalization within photoreceptor IS and cell bodies [18]. These observations suggested that MYO1C may participate in rhodopsin trafficking and that disruption of this process could contribute to photoreceptor dysfunction [17,18]. However, the molecular mechanism underlying this phenotype and whether MYO1C acts directly within photoreceptors remained unknown. Additionally, it was unclear whether MYO1C interacts with rhodopsin, which specific region of MYO1C mediates such a protein-protein interaction, and whether MYO1C function is required preferentially in rod or cone photoreceptors [18,19].

In the present study, we used a complementary combination of in-silico computer based analysis of protein-protein interaction, biochemical, cellular, imaging, and genetic approaches to investigate the role of MYO1C in rhodopsin trafficking and photoreceptor cell function. Protein-protein docking and biochemical interaction assays were used to examine potential molecular associations between MYO1C and rhodopsin and to assess the contribution of the MYO1C C-terminal cargo binding region. Live-cell imaging and fluorescence recovery after photobleaching (FRAP) were used to evaluate rhodopsin trafficking and membrane dynamics in cultured hTERT-RPE1 cells, while immunolabeling and high-resolution imaging approaches were used to examine the subcellular localization of MYO1C and rhodopsin in isolated murine rod photoreceptors. To define the physiological role of MYO1C in vivo, we evaluated retinal structure and function in global *Myo1c*-deficient mice across different ages and assessed rhodopsin distribution and mislocalized opsin protein accumulation in isolated photoreceptors and whole retina, respectively. Finally, to distinguish cell-autonomous effects and determine the relative contribution of MYO1C in individual photoreceptor subtypes, we generated and characterized rod-specific and cone-specific conditional *Myo1c* knockout mice. Together, these complementary approaches provide a framework for defining the molecular, cellular, and physiological functions of the myosin motor protein MYO1C in rhodopsin trafficking and for photoreceptor and retinal homeostasis.

## Materials and Methods

### Materials

All chemicals unless stated otherwise were purchased from Sigma-Aldrich (St. Louis, MO, USA) and were of molecular or cell culture grade quality. Cell culture reagents including Fetal Bovine Serum (FBS) were purchased from Gibco/ThermoFisher (Waltham, MA, USA).

### In Silico Modeling and Docking analysis

The complete structures of human and mouse MYO1C and Rhodopsin were obtained from AlphaFold (https://alphafold.ebi.ac.uk/), and the human protein sequences MYO1C (UNIPROT ID O00159) and RHO (UNIPROT ID P08100), using the available crystal structures of the human MYO1C N-terminus (PDB 4BYF), the mouse MYO1C C-terminus (PDB 4R8G), and the bovine RHO complete structure (1HZX) [19]. Using MEMDOCK (PMID: 27153621), the interaction between human MYO1C and RHO was predicted by protein-protein docking. The interaction site was visualized using the PyMOL Molecular Graphics System, Version 2.0, Schrödinger, LLC. The docking results were reconfirmed using another docking server, HADDOCK 2.4 [20].

### Construction of turboGFP-human rhodopsin and mCherry-human MYO1C recombinant proteins in pCDNA 3.1 plasmid

Human cDNA plasmids were obtained for turboGFP-human rhodopsin (NM_000539; Origene Clone#RG211328) and mCherry-human MYO1C (NM_001080779.1; GenecCopoeia Clone#EX-Z4322-M55) and cloned into the pCDNA3.1 vector. Sequencing, western blotting, and immunofluorescence imaging confirmed the proper construct generation, expression, and folding for a functional recombinant MYO1C protein interacting with F-actin.

### Fluorescence Recovery after Photobleach (FRAP) in COS1 cells

COS1 cells were transfected with mCherry-MYO1C pcDNA3.1 plasmid (mCherry-MYO1C), mCherry-ΔC-MYO1C pcDNA3.1 plasmid and turboGFP-rhodopsin pcDNA3.1 plasmid (GFP-rhodopsin). FRAP experiments were performed using an Olympus FluoView IX2 Inverted Confocal with FLIM Detector equipped with multiple lasers [21–24]. The experiments were performed using a 60X objective lens. The laser power was maintained at 100% for capturing photobleached regions of interest (ROI), and maintained at 10-20% for capturing recovery images Approximately 24 h post-transfection, using green (GFP) and red (mCherry) channels, an ∼2 μm ROI roughly close to the plasma membrane-cytoplasm interface with a visible distribution of GFP and mCherry signals was photobleached. In addition, an ∼2 μm region of reference was annotated as background ROI. Photobleaching was performed with a 488 nm laser for 30 sec and recovery was monitored using the same wavelength, for the next 2 min at an interval of ∼1 sec each. The acquired images were analyzed using Image J (NIH). Quantitative FRAP calculations were performed using published protocols and FRAP recovery values were normalized and plotted in the GraphPad Prism (version 9.3. San Diego, CA, United States) [21–24]. Fraction was calculated by dividing with normalized prebleach intensity values and base correcting with the lowest after-bleach intensity value. FRAP experiments were repeated thrice, using new transfections. Data was collected from three separate transfections and treatments and approximately 100-120 cells/condition were analyzed.

### Live cell rhodopsin trafficking in COS1 and hTERT-RPE1 cells

COS1 cells were transfected with mCherry-WT-MYO1C or mCherry-ΔC-MYO1C, and turboGFP-rhodopsin pcDNA3.1 plasmids, and then used to image rhodopsin trafficking [24]. As rhodopsin oligomerizes, it is possible to track its MYO1C-dependent movement. Post transfection, the cells on the glass bottom dish were scanned for transfections and suitable cells co-expressing both rhodopsin and MYO1C were identified for the imaging analysis. The laser was adjusted to ∼20% to achieve a sufficient signal and minimize photobleaching. Images were captured with an approximate interval of 1.1 sec for 1 or 3 min, n=∼10 cells with 1 or 3 min time frames were captured. Post-acquisition, the images were processed with Olympus FluoView viewer and Image J software (version 1.5), scaled, calibrated, and fed into a manual tracking plugin. The tracking data was plotted and velocity significance was analyzed by *t*-test in GraphPad Prism.

### Live-Cell Imaging of Rhodopsin-GFP Trafficking in hTERT-RPE1 Cells

To investigate the role of the MYO1C C-terminal domain in rhodopsin trafficking and membrane tethering, hTERT-RPE1 cells were transiently transfected with recombinant plasmids encoding mCherry-WT-MYO1C or a C-terminal deletion mutant of MYO1C (ΔC-MYO1C), together with wild-type GFP-Rhodopsin. This experimental design enabled comparison of rhodopsin trafficking in the presence of full-length MYO1C versus MYO1C lacking its C-terminal domain. Following transfection, cells were cultured for approximately 24 h in FluoroBrite™ DMEM supplemented with 5% FBS before live-cell imaging. Live-cell fluorescence imaging was performed using a Nikon AXR confocal microscope equipped with a 60X objective. Images were acquired using the NSPARC(SR) detector in Galvano unidirectional scanning mode with a dwell time of 1.6 μs. Rhodopsin-GFP and mCherry-MYO1C fluorescence was excited using 488 nm and 568 nm laser lines, respectively. Time-lapse images were acquired at 4-7 s intervals for approximately 5 min to monitor the dynamic trafficking and membrane localization of Rhodopsin-GFP. Image sequences were analyzed using ImageJ (NIH). Gaussian blur was applied to reduce image noise, and the Bleach Correction-Simple Ratio method was used to correct for photobleaching during time-lapse acquisition. Images were cropped and bilinearly resized to 2400 X 2400 pixels for subsequent visualization and analysis.

### Murine Photoreceptor Isolation and Immunostaining

Mice were euthanized by CO_2_ asphyxiation and cervical dislocation. The eyes were enucleated, and retinal tissue from the eye cups was surgically removed, and washed with 1XPBS. Retinal tissue was pre-incubated with Trypsin (0.05%) and Papain (5 U/mL) in a 3 mM N-acetyl-L-cysteine PBS buffer at 37°C for 15 minutes. Retinal tissue was then transferred to 100 μL of the enzyme mixture and incubate at room temperature for 5 minutes. After digestion, the enzymatic reaction was terminated with 100 µL of 10% FBS-DMEM, and 100 µL of the photoreceptor-retinal mixture was transferred to microscope slides and spread uniformly. The smear was fixed with 4% PFA or ice-cold methanol at room temperature for 5-10 minutes, and then processed for immunostaining [25].

### Mice

Twelve-week-old wild-type (WT) mice (C57BL/6J) were purchased from The Jackson Laboratories (RRID:IMSR_JAX:000664). All animals used in this study were genotyped and found to be negative for the known *Rd8* and *Rd1* mutations, as previously described by us (Table 1) [18,26]. Mice were provided a regular chow diet and water *ad libitum* and maintained at 24°C in a 12:12 hour light-dark cycle. All animal experiments were approved by the Institutional Animal Care and Use Committee of the University of Minnesota (IACUC protocol #38982A) and performed in compliance with the ARVO Statement for the use of Animals in Ophthalmic and Vision Research.

### Confocal Imaging

Mice were euthanized by CO_2_ asphyxiation and cervical dislocation. The eyes were enucleated, fixed with 4% PFA in 1X PBS, and embedded and paraffin sectioned. The tissue sections were processed and permeabilized with 0.5% TritonX-100 in 1X PBS at 4°C for 20 min. The tissue was blocked with 5% Bovine Serum Albumin in 1X PBS at room temperature for 1 hour. The primary antibody Rhodopsin 1D4 and anti-Myo1c M2

(dilution 1:100) was diluted in blocking solution and incubated at 4°C overnight. The tissue was washed with 0.1% TritonX-100 in 1X PBS at 4°C for 5 min, three times, and incubated for 1 hour at room temperature with secondary antirabbit and anti-Mouse IgG Alexa fluor conjugated (dilution 1:200) (Catalog no: A10524, Thermo Fisher Scientific, Waltham, MA, USA) and 1X Alexa Fluor 488 phalloidin (Catalog no: A12379, Thermo Fisher Scientific) (for f-actin labeling) in blocking solution. The tissue section was mounted on glass slides with VECTASHIELD® Antifade Mounting Medium with DAPI (Catalog no: H-1200-10, Vector Laboratories, Inc., Newark, CA, USA). Volumetric images were captured with a 60X objective and 0.1 μm Z-scan using a Next Generation Super-Resolution Nikon AX/AX R NSPARC Confocal microscope (Nikon Instruments Inc., Melville, NY, USA) at the University of Minnesota facility. The volumetric images were further analyzed using ImageJ or Fiji (ver 1.54f, NIH, USA).

### Generation of *Myo1c* knockout line

We previously generated *Myo1c* transgenic mice (*Myo1c^fl/fl^*) in C57BL/6N-derived embryonic stem cells, flanking exons 5 to 13 of the mouse *Myo1c* gene, which allowed us to specifically delete all *Myo1c* isoforms in a cell-specific manner [18]. Here, a complete *Myo1c*-knockout was generated by crossing *Myo1c^fl/fl^* mice with an F-actin Cre mouse strain (B6N.FVB-Tmem163Tg(ACTB-cre)2Mrt/CjDswJ) obtained from Jackson Labs. We refer to the *Myo1c^fl/fl^* x F-actin-Cre+ cross as *Myo1c* knockout (*Myo1c*-KO) mice [18].

### Generation of Rod Photoreceptor conditional *Myo1c*-KO mice

To obtain rod-specific *Myo1c*-KO mice, we crossed *Myo1c^flox/flox^* mice with the rod specific Rhodopsin-Cre+ mice (B6.Cg-*Pde6b^+^* Tg(Rho-icre)1Ck/Boc; JAX#015850; RRID:IMSR_JAX:015850) and obtained heterozygous *Myo1c^fl/wt^;*Rhodopsin-Cre+ mice. Primer sequences are listed in Table 1. Heterozygous *Myo1c^flox/wt^;*Rhodopsin-Cre+ mice were further mated with *Myo1c^fl/fl^* mice and litters were genotyped to obtain the experimental *Myo1c^flox/flox^;*Rhodopsin-Cre+ (*Myo1c;*Rho-Cre+) and littermate controls . All breeding pairs were genotyped and are negative for the *rd1* and *rd8* mutations.

### Generation of Cone Photoreceptor conditional *Myo1c*-KO mice

STOCK Tg(OPN1LW-cre)4Yzl/J Strain #:032911, RRID: IMSR_JAX:032911. HRGP-Cre was purchased from the Jackson Laboratory. The genetically modified strain expresses the Cre enzyme under the control of the human OPN1LW opsin promoter. Expression is cone photoreceptor specific. The HRGP-Cre+ FVB mouse was crossed with a *Myo1c^flox/flox^* mouse. Positive pups at day 21 were confirmed by PCR-based agarose gel genotyping. Primer sequences are listed in Table 1. F1 generation heterozygous mice were backcrossed with *Myo1c^flox/flox^* mice, and the subsequent generation mice were screened and selected for positive homozygous *Myo1c^flox/flox^* and HRGP Cre+ and negative for *Rd1* and *Rd8* mutations. Cone photoreceptor-specific Cre expression deleted *Myo1c* in cone photoreceptors to generate *Myo1c*;HGRP-Cre+ mice.

### Morphometry Analysis by Light Microscopy

The lengths of the photoreceptor OS in WT and global and conditional *Myo1c*-KO animals, from H&E sections of retinas, were imaged (Keyence BZ-X800 microscope) and measured at 8 consecutive points (at 100 μm distances) from the optic nerve (ON) [18,24]. The OS length was measured from the base of the OS to the inner side of the retinal pigment epithelium [18,24]. The total number of layers of nuclei in the ONL of retinal sections through the optic nerve (ON) was imaged (Keyence BZ-X800 microscope) and measured at 8 locations around the retina, four each in the superior and inferior hemispheres, starting at 100 μm from the ON. Retinal sections (*n* = 5-7 retinal sections per eye) from *n* = 8 mice for each genotype and timepoint were analysed. Two-way ANOVA with Bonferroni post-tests compared *Myo1c*-KO to WT mice at each segment measured.

### Immunoblot Analysis

Total protein from cells or mouse tissues (*n* = 3 per genotype) were extracted using the M-PER protein lysis buffer (ThermoScientific, Beverly, MA) containing protease inhibitors (Roche, Indianapolis, IN) [27,28]. Approximately 25 μg of total protein was electrophoresed on 4-12% SDS-PAGE gels and transferred to PVDF membranes. Membranes were probed with primary antibodies against anti-*Myo1c* (1:250), 1D4-rhodopsin (1:1000, Sigma), R/G cone opsins (1:1000, Sigma), mCherry (1:1000, Sigma), and β-Actin or Gapdh (1:10,000, Sigma) in antibody buffer (0.2% Triton X-100, 2% BSA, 1X PBS) [45,59]. HRP-conjugated secondary antibodies (BioRad, Hercules, CA) were used at 1:10,000 dilution. Protein expression was detected using a LI-COR Odyssey system, and relative intensities of each band were quantified (densitometry) using Image *J* software version 1.49 and normalized to their respective loading controls. Each Western blot analysis was repeated thrice.

### Scotopic Electroretinography (ERG) Tests

For scotopic ERG measurements, the grouped mice were dark-adapted overnight and then anesthetized individually in an induction chamber with ∼4% isoflurane mixed with ∼25% oxygen and ∼75% nitrogen with a constant flow of 1 L/min. As soon as the mouse was sedated, the isoflurane was reduced to ∼1-2% with continuous oxygen and nitrogen flow rate. The mouse was kept on the Celeris ERG measuring heated platform, and ∼1-2% isoflurane was supplied with the nose cone, and pupils were dilated with Tropicamide 1%, (Sandoz #61314035502) and Phenylephrine 2.5% Eye Drops (AKORN INC #17478020115). A drop of 2.5 percent Hypromellose ophthalmic solution (Systane â Alcon) was spread onto the cornea, and the Celeris Light Guide Electrodes (LGE) electrodes were placed on the cornea with an impedance below 10kΩ. The scotopic ERG electrical responses were measured from 0.01 to 10 cd.s/m^2^ range of impulse flash intensities. The recorded signals were processed with the Celeris Diagnosys software to export raw files. The same mouse was kept under anesthesia for rod response recovery after bleaching; the rod response recovery after bleaching Celeris ERG protocol was used with a pulse frequency of 1 and pulse intensity of 1 cd.s/m^2^. The protocol was set to acquire pre-bleach responses and initiate the rod photoreceptor bleaching using a high-intensity continuous light impulse for 180 secs. After bleaching, the responses were recorded, and amplitudes of the *a*- and *b*-waves were measured under scotopic conditions every minute for 10 min post-bleaching. The time-dependent ERG recovery responses were plotted in GraphPad Prism, and curve fitting with nonlinear regression one phase association was performed on the recovered a-wave amplitudes to get the half-life, which suggests the time taken to recover half of the total ERG response recovery measured, using the formula Y=Y0 + (Plateau-Y0)*(1-exp(-K*x)). The equation is explained on the GraphPad website (https://www.graphpad.com/guides/prism/latest/curve-fitting/reg_exponential_association.htm). After the scotopic ERG protocols, the mouse was monitored for recovery from anesthesia. The responses were recorded and amplitudes were measured and plotted in GraphPad Prism [19,24,27,28].

### Optical Coherence Tomography and Fundus Imaging

The mice from various groups were sedated with 5% isoflurane with adjustable room airflow using a Kent Scientific mouse anesthesia machine. After sedation, the isoflurane was reduced to 2%, and the mouse eyes were dilated with a 1:1 mix of Tropicamide 1.0% Ophthalmic Solution (Cat. No. NDC 61314-355-02 Sandoz) and Phenylephrine HCl 2.5% Solution (Cat. No. NDC 17478-201-15 Akorn). Alcon Systane Lubricant Eye Gel was applied to moisturize the eye, and the Phoenix Micron IV image-guided OCT2 instrument, equipped with a specialized objective lens, was used to capture OCT images.

### Light-adapted Photopic ERG

Light adapted mice were sedated and eyes were dilated, as outlined in dark-adapted mice method section. To measure the photoreceptor cone response *a*-wave, *b*-wave, and retinal ganglion response photopic ERG, Celeris ERG protocol was performed under light adapted condition with various pulse frequencies and color wavelengths. The amplitudes were recorded and plotted in GraphPad Prism.

### Immunohistochemistry and Fluorescence Imaging

Mice were euthanized by CO_2_ asphyxiation and cervical dislocation. Eyes were enucleated and fixed with either 4% PFA in 1X PBS or in Davidson’s fixative for 4 h at 4^0^C. Paraffin-embedded retinal sections (∼10 μm) were processed for antigen retrieval and immunofluorescence. Primary antibodies were diluted in blocking solution as follows: anti-MYO1C M2 (1:100) [18], anti-rhodopsin 1D4 for mouse rod opsin (1:500; Millipore, St. Louis, MO, USA), and 4′,6-diamidino-2-phenylendole (DAPI; 1:5000, Invitrogen) or Hoechst (1:10,000, Invitrogen) to label nuclei. All secondary antibodies (Alexa Fluor 488 or Alexa Fluor 594) were used at 1:5000 concentrations (Molecular Probes, Eugene, OR, USA). Optical sections were obtained with a Leica SP8 confocal microscope (Leica, Wetzlar, Germany) and processed with the Leica Viewer software or using a Keyence BZ-X800 microscope. After mounting, images were captured using 40X and 60X objectives. All fluorescently labeled retinal sections on slides were analyzed using ZEISS ZEN 3.4 software package, Image J or Fiji (NIH), and intensities were quantified, and data were plotted in GraphPad Prism. Immunostaining was quantified using ImageJ. Briefly, the image was opened using ImageJ software and free hand tool was used to draw the region of interest in the retinal layers. From Analyze menu, measurements were set to calculate area-integrated intensity and mean gray value. The background values were noted in areas with no fluorescence. Corrected fluorescence was measured as difference of integrated density and area, adjusted for background fluorescence. The quantified values from the multiple ROIs were plotted and averaged. Statistical significance was measured using two-tailed *t*-test (gaussian distribution, unpaired). Significance was considered as having p-value <0.05. Statistical Analysis was done using Graph Pad Prism.

### Purification of Rhodopsin and Absorbance Spectroscopy

Mice were dark-adapted for 12-16 hours. Under single source red light, the mice were then (CO2) euthanized, and the retina was harvested surgically. In strict dark conditions, the retinal tissues were homogenized with 20 mM bis-tris propane, 150 mM NaCl, and 1 mM EDTA buffer pH 7.5 with protease inhibitor. The homogenates were centrifuged 15 min at 16,000g refrigerated. The supernatants were discarded, and the pellets solubilized for 1 hour on a rotating platform at 4^0^C in 20 mM bis-tris propane, 150 mM NaCl, 20 mM n-dodecyl-b-D-maltoside (DDM) buffer pH 7.5 with protease inhibitor. The lysate was centrifuged for 1 hour at 16,000g at 4^0^C. The supernatant was incubated for 1 hour with 30 µL HighSpec Rho1D4 MagBeads (Cat 33299 Cube Biotech Germany). The resin was washed on a magnetic stand with 20 mM bis-tris propane, 500 mM NaCl pH7.5 buffer two times and three times with low salt 20 mM BTP, 100 NaCl, and 2 mM DDM buffer. The VAPA peptides dissolved in low salt buffer were used at a concentration of 0.1mg/mL volume 60 µL to elute the rod opsins from 1D4 resin. The eluted Opsin was analyzed on an Agilent Cary 60 spectrophotometer Instrument; the measured absorbances were plotted in GraphPad Prism Version 10.1 and calculated for free Opsin using a 280/ 500 nm absorbance ratio, using the extinction coefficient ε_500_ = 40,600 M^−1^ cm^−1^. The concentration of ligand-free opsin was calculated using the extinction coefficient ε_280_ = 81,200 M^−1^ cm^−1^ [28].

### Statistical Analysis

Data is expressed as means ± standard error mean, statistical analysis by ANOVA and student *t*-test. Differences between means were assessed by Tukey’s honestly significant difference (HSD) test. P-values below 0.05 (p<0.05) were considered statistically significant. For western blot analysis, relative intensities of each band were quantified (densitometry) using the Image J software version 1.54 and normalized to Ponceau S stain. Statistical analysis was carried out using GraphPad Prism v 10.1.

## RESULTS

### In silico protein docking analysis identifies a putative MYO1C-rhodopsin binding region

The complete structure of human MYO1C and human rhodopsin was obtained from the human protein sequences MYO1C (UNIPROT ID O00159) and RHODOPSIN (UNIPROT ID P08100) against the available crystal structure of human MYO1C N’-terminus PDB 4BYF and mouse MYO1C C’-terminus PDB 4R8G and bovine Rhodopsin complete structure 1HZX. Using HADDOCK 2.4, the interactions between human MYO1C and RHODOPSIN were predicted by protein-protein docking. This analysis suggested a possible interaction between MYO1C and rhodopsin at their respective C’-terminus domains, where rhodopsin has a conserved VxPx motif that is essential for its trafficking from the photoreceptor IS to the OS (**Figure 1**). Interestingly, C-terminus rhodopsin mutations that cause rhodopsin mislocalization in RP [1–4], fall within the putative MYO1C-rhodopsin interaction domain and are predicted to affect MYO1C-rhodopsin binding and subsequent trafficking (**Figure 1**).

**Figure 1.**
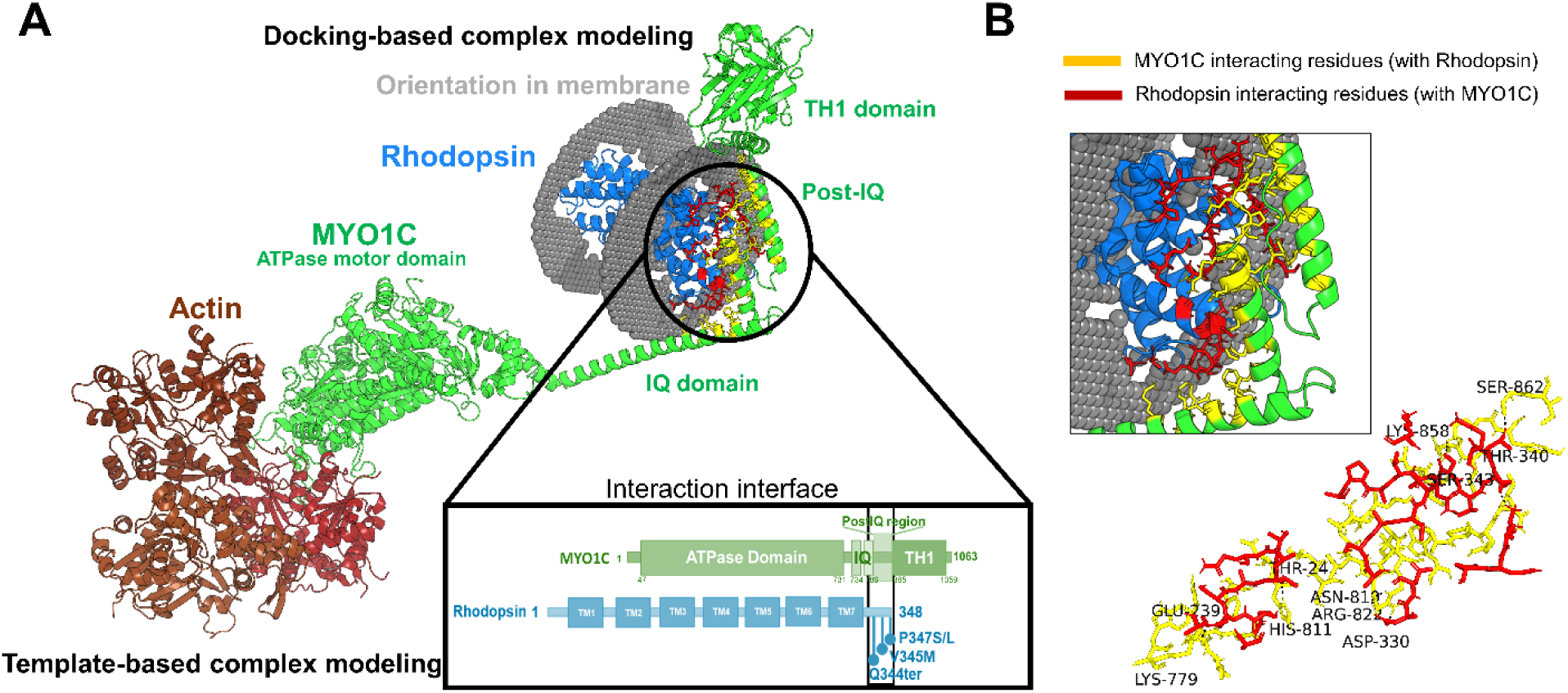
Virtual analysis of the putative binding interface between MYO1C and rhodopsin. (**A**) Computational modeling identified a potential interaction interface between MYO1C and rhodopsin, and MYO1C ATPase domain and F-Actin. The predicted binding region is highlighted, and the major domains of MYO1C-rhodopsin are indicated by a black circle. The C-terminal cargo-binding domain of MYO1C is predicted to interact with the rhodopsin C-terminal VxPx ciliary targeting motif (black box), a critical trafficking signal and mutational hotspot associated with rhodopsin mislocalization and the development of retinitis pigmentosa (RP). (**B**) Amino acid residues within the putative MYO1C-rhodopsin interaction interface are shown in yellow for MYO1C and red for rhodopsin.

### Generation of tagged full-length MYO1C, full-length Rhodopsin, and MYO1C C-terminus deletion constructs

The IQ domain of motor proteins (within the C’-terminus) is predicted to be important for cargo binding and trafficking [17,24] . In silico docking analysis suggested that either the IQ2 or the post-IQ domain of MYO1C interacts with the C’-terminus of rhodopsin (**Figure 1**). To test this prediction, we generated mCherry-tagged full-length MYO1C (mCherry-WT-MYO1C), GFP-tagged full-length rhodopsin (GFP-Rhodopsin), and mCherry-tagged MYO1C C-terminus deletion construct (ΔC-MYO1C) separately in the pCDNA3.1 plasmid (**Supplementary Figure S1**). Constructs and their expression were confirmed respectively by Sanger sequencing and by western blotting using the mCherry (for MYO1C expression) or GFP (for Rhodopsin expression) antibody (**Supplementary Figures S1A, S1B**).

### The C-terminal region of MYO1C is important for binding to Rhodopsin

COS1 cells were co-transfected with (a) mCherry-WT-MYO1C and GFP-Rhodopsin, (b) GFP-Rhodopsin alone, or (c) mCherry-ΔC-MYO1C (C-terminal deletion mutant) and GFP-Rhodopsin. Anti-GFP antibody was used for pull-down assays, and anti-mCherry antibody was used for Western blot analysis. Immunoblotting revealed robust co-immunoprecipitation of full-length WT-MYO1C with full-length Rhodopsin (**Supplementary Figure S1C**). In contrast, no interaction was detected between ΔC-MYO1C and Rhodopsin, indicating that the C-terminal region of MYO1C is required for binding to Rhodopsin (**Supplementary Figure S1C**). These findings suggest that the C-terminal cargo-binding region of MYO1C, which contains the IQ and post-IQ domains, plays a critical role in mediating the interaction between MYO1C and Rhodopsin (**Supplementary Figures S1**).

### The C’-terminus region of MYO1C is required for Rhodopsin trafficking in COS1 cells and to the cilium in hTERT-RPE1 cells

To further confirm that the C-terminal region of MYO1C mediates its interaction with and trafficking of rhodopsin, hTERT-RPE1 (immortalized human retinal pigment epithelial) cells were co-transfected with either (a) mCherry-WT-MYO1C and GFP-Rhodopsin or with (b) mCherry-ΔC-MYO1C (C-terminal deletion mutant) and GFP-Rhodopsin. Live-cell trafficking of GFP-Rhodopsin-positive vesicular foci was monitored by confocal microscopy (**Figure 2A**). Data from hTERT-RPE1 cells demonstrated efficient trafficking of GFP-Rhodopsin in the presence of wild-type MYO1C (**Figures 2B-2D**). In contrast, deletion of the MYO1C C-terminal domain (ΔC-MYO1C) markedly impaired GFP-Rhodopsin trafficking in both cell types (**Figures 2A-2D**), indicating that the C-terminal region of MYO1C is required for proper rhodopsin trafficking.

**Figure 2:**
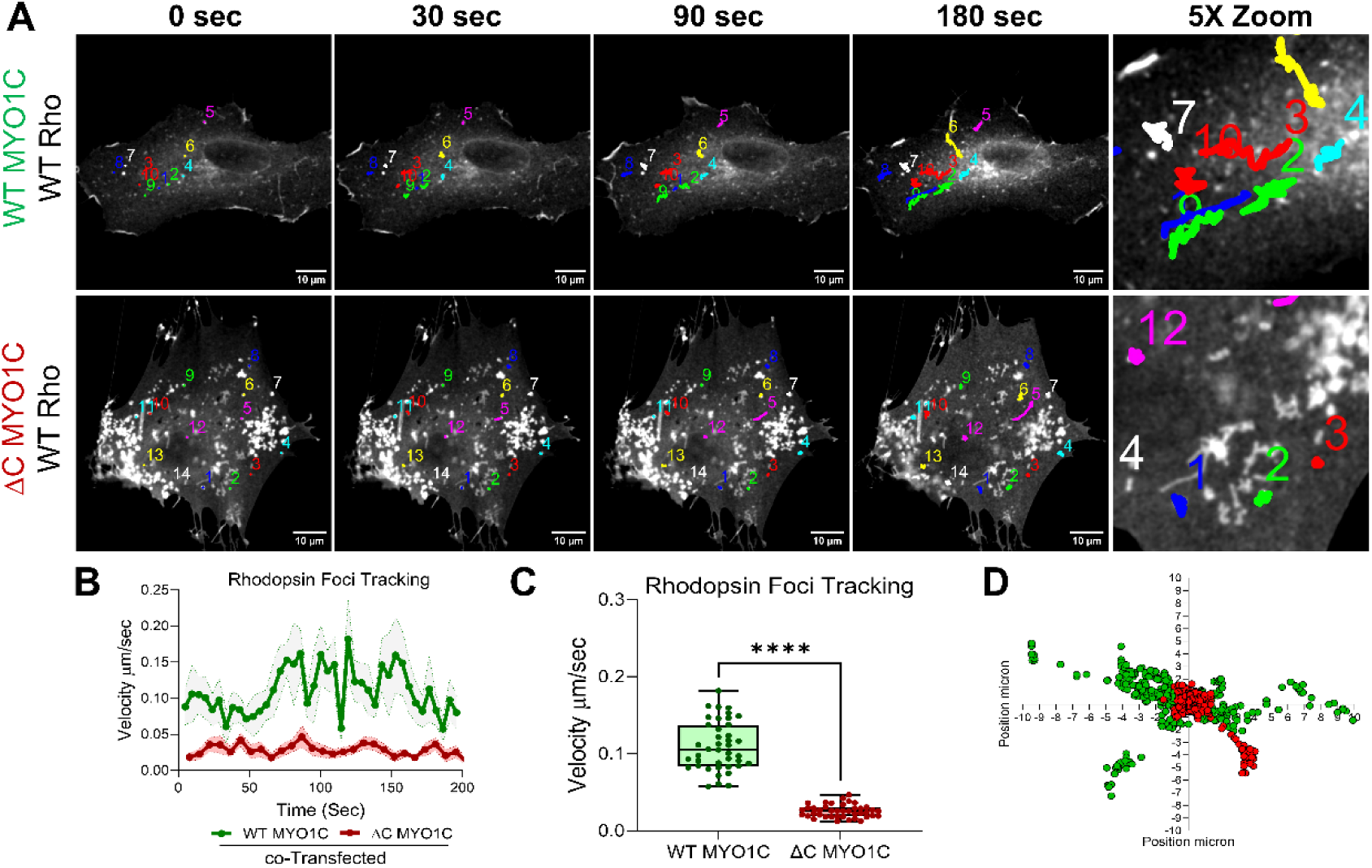
Live-cell imaging of GFP-rhodopsin foci trafficking in COS1 and hTERT-RPE1 cells. (**A**) hTERT-RPE1 cells were co-transfected with mCherry-WT-MYO1C and GFP-WT-rhodopsin or with mCherry-ΔC-MYO1C and GFP-WT-rhodopsin. hTERT-RPE1 cells expressing both GFP and mCherry were used in our analysis. Individual GFP-rhodopsin foci (color-coded/numbered) tracking and movement data was obtained from 0-120 sec. (**B-D**) Defective GFP-rhodopsin trafficking was observed in cells expressing C-terminus deleted MYO1C (ΔC-MYO1C). Statistical significance was determined using ANOVA. \*\*\*\**P* < 0.001.

### Loss of the MYO1C C-Terminal Domain disrupts Rhodopsin Trafficking and Membrane Localization in hTERT-RPE1 cells

To investigate the role of MYO1C in rhodopsin trafficking and membrane tethering, hTERT-RPE1 cells were co-transfected with full-length GFP-Rhodopsin and either full-length mCherry-MYO1C or a C-terminal deletion mutant of mCherry-MYO1C. Live-cell imaging was performed by acquiring Rhodopsin-GFP and mCherry-MYO1C fluorescence images at 4-7 s intervals for approximately 5 min. In cells expressing full-length MYO1C, time-lapse imaging revealed dynamic translocation of MYO1C from the cytoplasm toward the plasma membrane, accompanied by enrichment and colocalization of Rhodopsin-GFP at membrane-associated structures (**Figure 3A**). These observations are consistent with a proposed role for MYO1C in facilitating the trafficking and sequential tethering of Rhodopsin at the plasma membrane. In contrast, deletion of the MYO1C C-terminal region markedly altered Rhodopsin trafficking. The C-terminal domain, which contributes to Rhodopsin binding and membrane interaction, was required for efficient redistribution of Rhodopsin-GFP to the plasma membrane. Cells expressing the C-terminal MYO1C deletion construct exhibited prominent cytoplasmic Rhodopsin-GFP aggregates or foci, with reduced membrane-associated Rhodopsin compared with cells expressing full-length MYO1C (**Figure 3B**). These findings suggest that loss of the MYO1C C-terminal domain disrupts the trafficking and membrane tethering of Rhodopsin, resulting in abnormal intracellular accumulation of the protein (**Figures 3A, 3B**). Together, these live-cell imaging data support a model in which MYO1C coordinates the trafficking and membrane tethering of Rhodopsin, thereby facilitating its proper incorporation into the plasma membrane. Disruption of the MYO1C C-terminal domain compromises this process and alters the normal membrane distribution and dynamics of rhodopsin.

**Figure 3.**
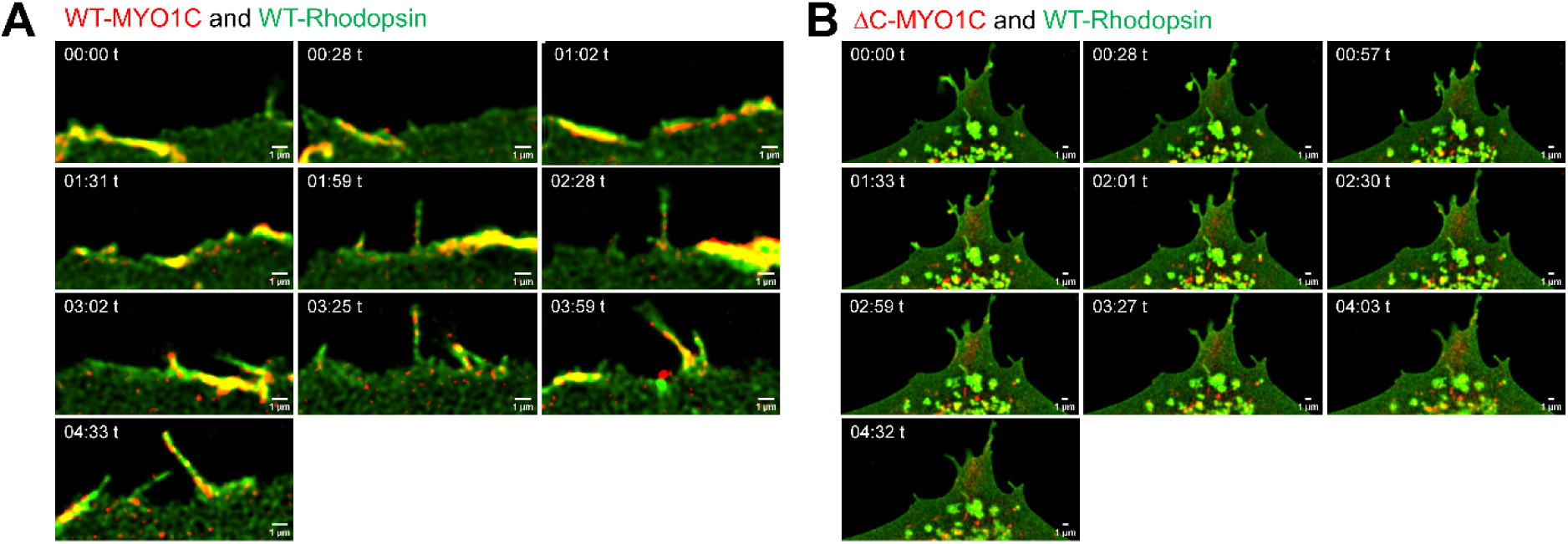
Loss of the MYO1C C-terminal domain disrupts Rhodopsin trafficking and membrane localization in hTERT-RPE1 cells. (**A**) Representative live-cell time-lapse images of hTERT-RPE1 cells co-expressing GFP-Rhodopsin and full-length mCherry-MYO1C. Full-length MYO1C dynamically redistributed from the cytoplasm toward the plasma membrane, accompanied by enrichment of GFP-Rhodopsin at membrane-associated structures. (**B**) Representative live-cell images of hTERT-RPE1 cells co-expressing GFP-Rhodopsin and a C-terminal deletion mutant of mCherry-MYO1C (ΔC-MYO1C). Loss of the MYO1C C-terminal region was associated with prominent intracellular GFP-Rhodopsin aggregates/foci and reduced membrane-associated Rhodopsin compared with cells expressing full-length MYO1C. Images were acquired at 4-7 s intervals for approximately 5 min. Scale bars, 1 μm.

### MYO1C is required for Rhodopsin trafficking and localization to the primary cilium in hTERT-RPE1 cells

We chose to study hTERT-RPE1 cells, because they have a cilium that protrudes from a pocket in the apical membrane. Transfected hTERT-RPE1 cells were serum starved to induce formation of cilia. In serum-starved hTERT-RPE1 cells expressing mCherry-WT-MYO1C, GFP-rhodopsin efficiently trafficked to the primary cilium and co-localized with WT-MYO1C and with acetylated α-tubulin in the cilium (**Supplementary Figure S2A**), which contrasted with the ΔC-MYO1C transfected hTERT-RPE1 cells, where rhodopsin trafficking to the primary cilium was impeded (**Supplementary Figure S2B**). Taken together, these findings demonstrate that MYO1C is associated with rhodopsin trafficking during ciliary targeting and further support an essential role for the MYO1C C-terminal domain in mediating rhodopsin trafficking and localization to the primary cilium.

### FRAP analysis of RHO-GFP recovery in hTERT-RPE1 cells

To study the movement of rhodopsin to the plasma membrane and its dependence on MYO1C, we developed a protocol based on FRAP using hTERT-RPE1 cells transfected with GFP-rhodopsin. For these experiments, an ∼2 μm circular region covering the plasma membrane was bleached, so that any fluorescence recovery must result from the movement/trafficking from the cytoplasm to the plasma membrane. **Figure 4A** shows a representative example of recovery of GFP-rhodopsin fluorescence in co-transfected WT-MYO1C expressing cells, after 10-60 sec of photobleaching (**Figures 4C, 4F, 4H**). In contrast, there was a significant delay in the recovery of GFP-rhodopsin fluorescence in cells co-transfected with ΔC-MYO1C (**Figures 4B, 4D, 4F, 4H**). The time taken to reach half of the fluorescence recovery (half time *t_1/2_*), as determined from exponential fit of the data, showed that in hTERT-RPE1 cells, ΔC-MYO1C transfected cells was a mean of 73.35, while in WT-MYO1C cells was a mean of 35.1 (**Supplementary Figure S3A**). The diffusion coefficient (D) is a quantitative measure of a molecule’s mobility within the cell, expressed in μm²/s. Rhodopsin exhibited a significantly lower diffusion coefficient (D) under conditions involving C-terminal deletion of MYO1C (ΔC-MYO1C co-transfected cells), further suggesting that the MYO1C C-terminus contributes to rhodopsin mobility and trafficking from the cytosol to the plasma membrane (**Figure 4H**). In contrast, MYO1C diffusion was not affected by deletion of its C-terminus, likely because its N-terminus retains actin-binding activity and functional ATPase activity required for the power stroke movement (**Figure 4G, Supplementary Figure S3B**). Overall, these FRAP studies demonstrate that loss of the MYO1C C-terminal region substantially impairs the dynamics of Rhodopsin delivery to the plasma membrane, providing functional evidence that the MYO1C C-terminus is important for efficient Rhodopsin trafficking.

**Figure 4.**
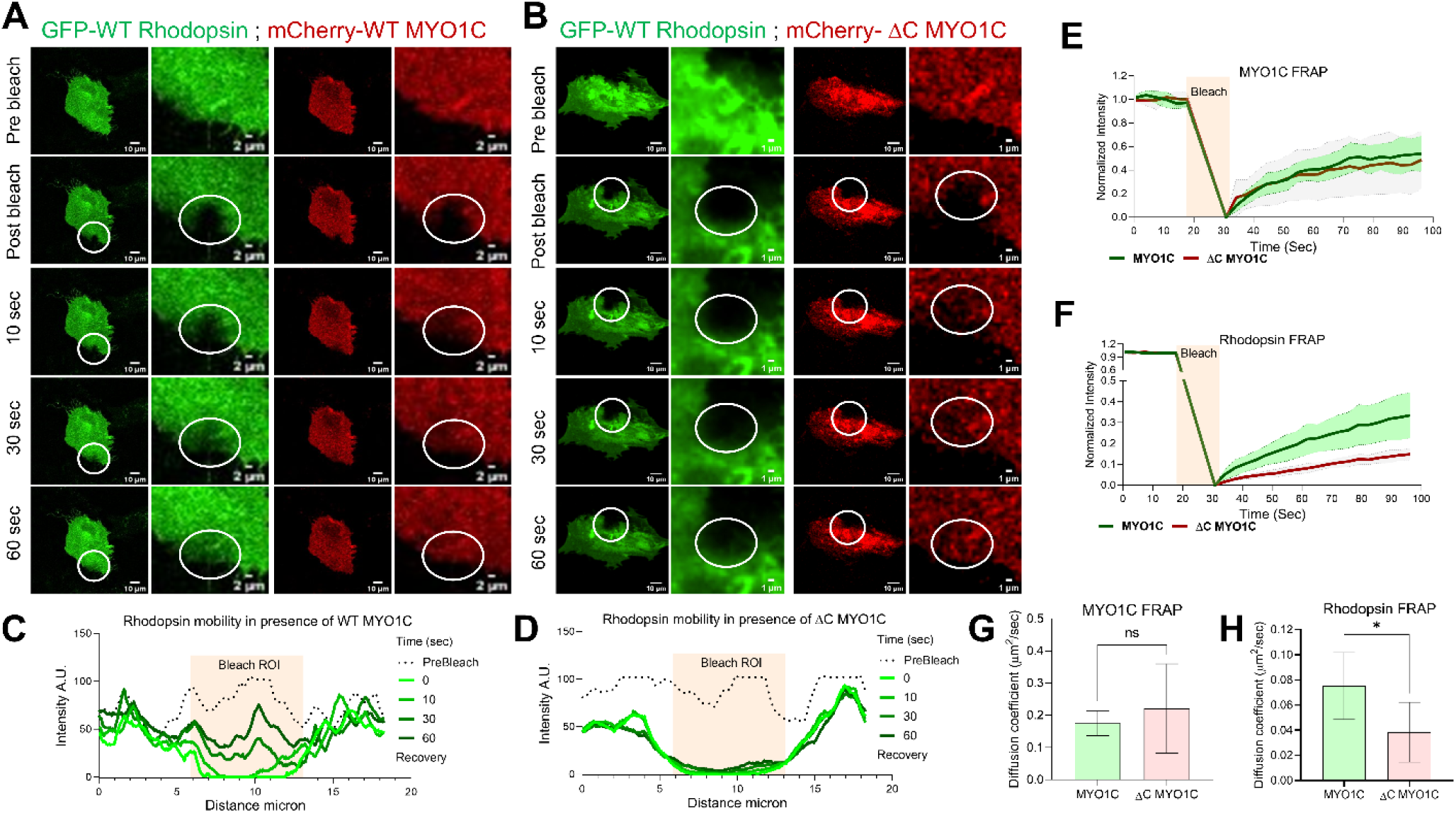
Loss of the MYO1C C-terminal domain impairs Rhodopsin mobility and recovery at the plasma membrane. hTERT-RPE1 cells expressing GFP-Rhodopsin together with either WT-MYO1C or ΔC-MYO1C were analyzed by fluorescence recovery after photobleaching (FRAP). A circular region encompassing the plasma membrane was photobleached, and fluorescence recovery was monitored over time. (**A, C**) Representative FRAP images and quantification of GFP-Rhodopsin in cells co-expressing WT-MYO1C, showing robust fluorescence recovery following photobleaching. (**B, D**) Representative FRAP images and quantification of GFP-Rhodopsin in cells co-expressing ΔC-MYO1C, showing markedly delayed fluorescence recovery. (**C, D**) Representative fluorescence recovery curves for GFP-Rhodopsin in WT-MYO1C- and ΔC-MYO1C-expressing cells, respectively. (**E, F**) Quantification of fluorescence recovery kinetics demonstrating reduced recovery and prolonged half-time of recovery (*t*1/2) for rhodopsin following deletion of the MYO1C C-terminal region. (**E, G**) Quantification of MYO1C diffusion coefficient showing no significant difference between WT-MYO1C and ΔC-MYO1C. (**H**) Quantification of the Rhodopsin diffusion coefficient demonstrating significantly reduced Rhodopsin mobility in ΔC-MYO1C-expressing cells compared with WT-MYO1C-expressing cells. Data are presented as mean ± SEM. Statistical significance was determined using ANOVA. \**P* < 0.05.

### MYO1C localizes to the Inner and Outer Segments of Isolated Murine Rod Photoreceptors

To determine the subcellular localization of MYO1C within rod photoreceptors, murine photoreceptors were isolated and subjected to immunofluorescence staining for MYO1C, acetylated tubulin, and rhodopsin. Acetylated α-tubulin was used to visualize the photoreceptor connecting cilium (CC), whereas rhodopsin (1D4) was used as a marker of the rod OS. Fluorescence imaging revealed MYO1C immunoreactivity in both the inner segment (IS) and outer segment (OS) compartments of isolated murine rod photoreceptors (**Figure 5**). MYO1C staining overlapped with the rhodopsin-positive OS and was also detected within the IS region. The merged images further demonstrated the spatial distribution of MYO1C relative to acetylated α-tubulin and rhodopsin, supporting the presence of MYO1C within distinct structural compartments of rod photoreceptors (**Figure 5**). The localization of MYO1C within the photoreceptor IS and OS suggests that MYO1C may contribute to the intracellular trafficking and membrane-associated organization of photoreceptor proteins. MYO1C presence in the OS, together with its association with rhodopsin-positive structures, is consistent with a potential role for MYO1C in rhodopsin trafficking and/or membrane tethering (**Figure 4**) within rod photoreceptors.

**Figure 5.**
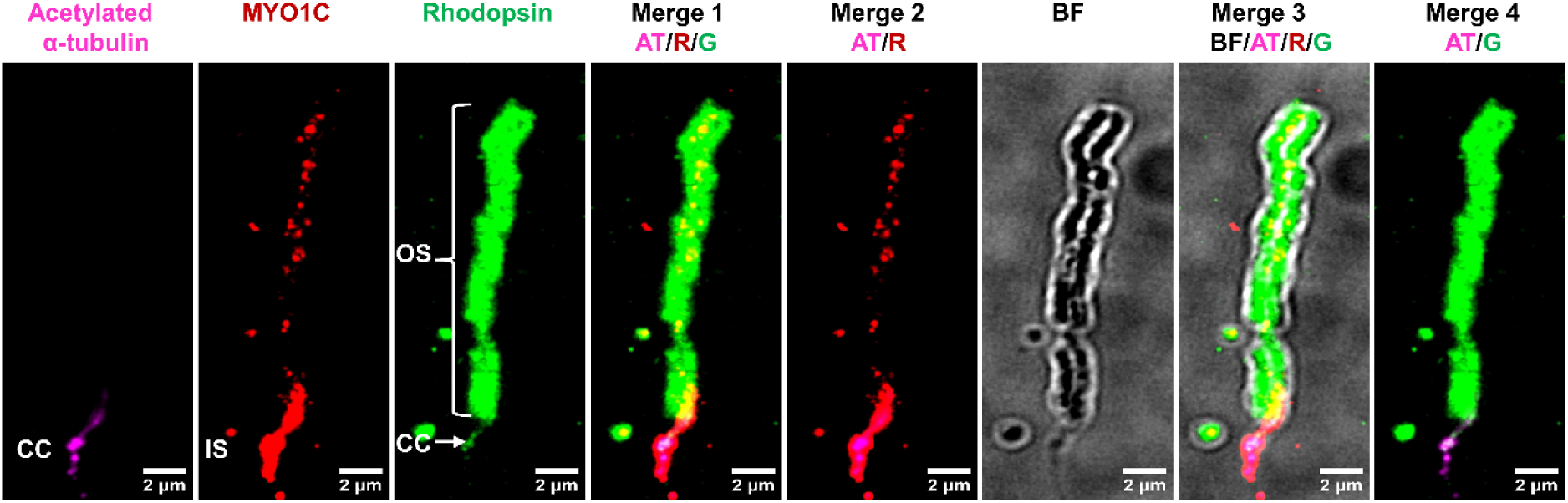
MYO1C localizes to the inner and outer segments of isolated murine rod photoreceptors. Isolated murine photoreceptors were immunostained for acetylated α-tubulin (purple), MYO1C (red), and Rhodopsin (green). Representative fluorescence images show the localization of MYO1C relative to the photoreceptor axoneme and Rhodopsin-positive outer segment. Merged images demonstrate MYO1C localization within the inner segment and outer segment compartments of rod photoreceptors. CC, connecting cilium, OS, outer segment, IS, inner segment, BF, bright field.

### Age-dependent retinal dysfunction in *Myo1c*-KO mice

We previously demonstrated that global *Myo1c*-KO mice exhibit severe mislocalization of rod opsin to the photoreceptor inner segment (IS) and cell bodies [18]. To determine whether this abnormal rhodopsin localization is associated with functional deficits, we performed a comprehensive, non-invasive assessment of retinal structure, vasculature, and visual function in *Myo1c*-KO mice, using optical coherence tomography (OCT), fluorescein angiography (FA), and scotopic and photopic electroretinography (ERG).

### *Myo1c*-KO mice display age-dependent decline in retinal function

Scotopic electroretinography (ERG) was performed to determine whether rhodopsin mislocalization observed in *Myo1c*-KO mice is associated with age-dependent retinal dysfunction. At 2-months of age, scotopic ERG responses were comparable between WT and *Myo1c*-KO mice, with no significant differences in a-wave and b-wave amplitudes (**Figures 6A-6C**). By 6-months of age, *Myo1c*-KO mice exhibited significant reductions in both scotopic a- and b-wave amplitudes, which persisted through 10-months of age (**Figures 6A-6C**). Photopic negative ERG responses showed greater variability, with some but not consistent age-dependent reduction in photopic responses between WT and *Myo1c*-KO mice (**Figures 6D-6F**). These findings indicate that loss of MYO1C results in progressive, predominantly rod-mediated retinal dysfunction, while cone function appears to be relatively preserved.

**Figure 6:**
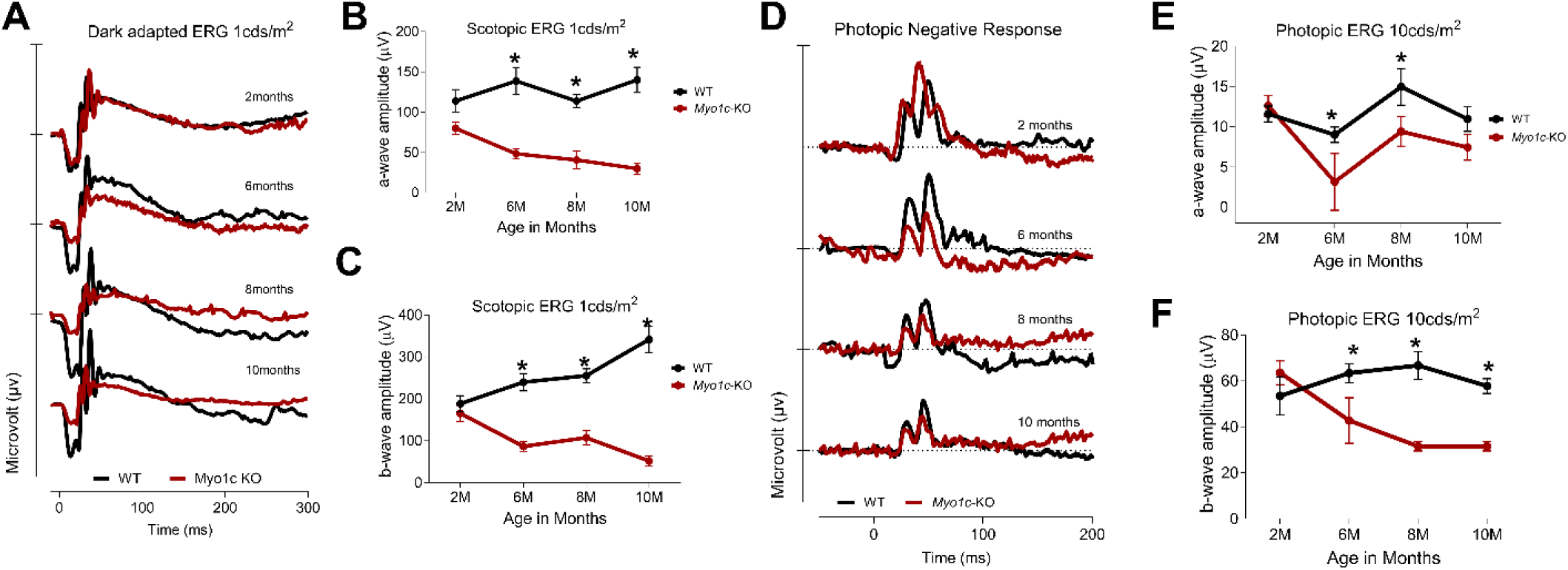
Genetic deletion of *Myo1c* in mice results in decreased visual function. (**A**) Dark-adapted scotopic ERGs were recorded in response to increasing light intensities in cohorts of control wild-type/WT (black-traces) and *Myo1c*-KO (red-traces) mice from 2-to 10-months of age. Two-months old *Myo1c*-KO mice had similar dark-adapted *a*- and *b*-wave amplitudes compared with controls (post-hoc ANOVA: *a*-waves, n.s. not significant.). (**B, C**) Starting at six-months *Myo1c*-knockout mice had lower dark-adapted *a*- and *b*-wave amplitudes compared with controls (post-hoc ANOVA: *a*-waves, \**P* < 0.05; *b*-waves, \**P* < 0.05). Photoreceptor cell responses (*a*-waves), which drive the *b*-waves, were equally affected in 6-months old *Myo1c*-KO animals (both reduced on average between 38-45% of WT animals). (**D-E**) Photopic negative responses (PhNRs) in *Myo1c*-KO mice were also significantly different from those of age-matched WT controls. Data are expressed as mean ± S.E. *Myo1c*-KO mice *and* WT mice, *n*=8 per genotype and age-group; 50:50 ratio of male and female.

### Rod photoreceptors in *Myo1c*-KO mice accumulate significant levels of apo-opsin

Because *Myo1c*-KO mice exhibit reduced visual responses, we hypothesized that this phenotype may be associated with disrupted opsin homeostasis and accumulation of unliganded (apo-) opsin in rod photoreceptors. To test this hypothesis, we quantified rhodopsin and apo-opsin levels in isolated retinal protein fractions from *Myo1c*-KO and WT mice using UV-visible spectrophotometry. Rhodopsin and apo-opsin levels were determined based on experimentally measured and theoretical 280/500-nm absorbance ratios. In dark-adapted 6-month-old mice, *Myo1c*-KO photoreceptors exhibited a significant (∼22%) increase in apo-opsin levels compared with age-matched WT controls (**Figures 7A; quantified in 7C**). This difference was further pronounced at 10-months of age, when apo-opsin levels were significantly (∼41%) higher in *Myo1c*-KO mice than in WT controls (**Figures 7B; quantified in 7C**). These findings demonstrate an age-dependent accumulation of unliganded and mistrafficked opsin in *Myo1c*-deficient rod photoreceptors. Accumulation of apo-opsin has previously been shown to produce constitutive activation of the phototransduction cascade in darkness. Such constitutive activity could effectively increase the background signal and reduce the dynamic range or gain of phototransduction. Thus, the age-dependent accumulation of unliganded opsin observed following loss of MYO1C may contribute to the impaired photoreceptor function and altered rod and cone phototransduction kinetics observed in *Myo1c*-KO mice.

**Figure 7:**
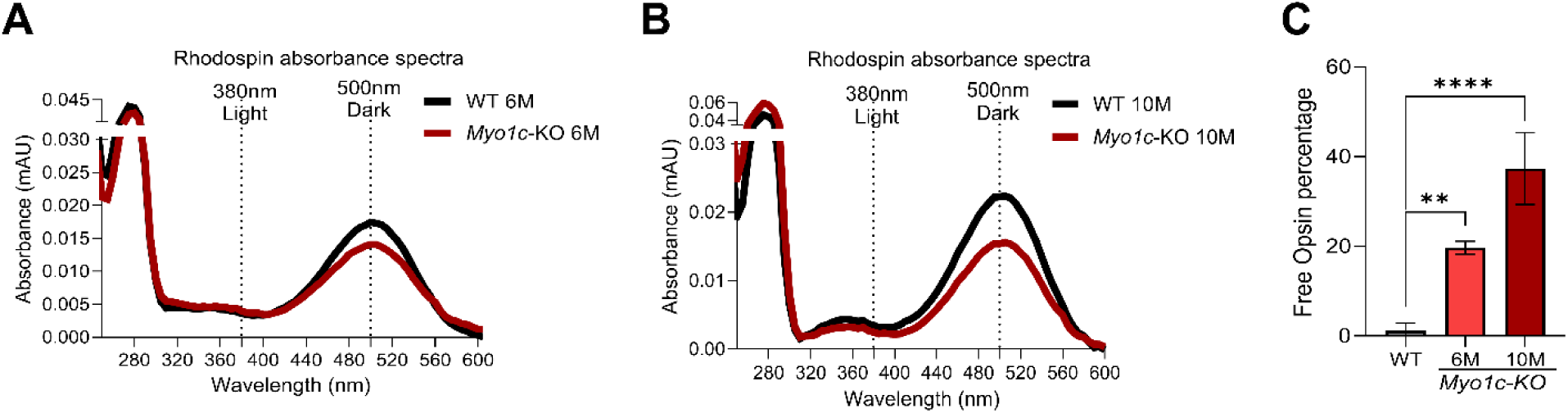
Presence of apoprotein opsin in rod photoreceptors of *Rbpr2^-/-^* mice. Rhodopsin absorbance spectra in dark-adapted (500 nm) animals at 6-months of age (**A**) and at 10-months of age (**B**), and threshold-based quantification of mislocalized Rhodopsin in the IS of retinas from WT and *Myo1c-KO* mice (**C**). Values are presented as ±SD. 2-way ANOVA, \*\**P* < 0.01; \*\*\*\**P* < 0.001. n=8-10 animals per group.

### Rationale for generating Rod- and Cone-Specific *Myo1c* Conditional Knockout Mice

Global *Myo1c* deficiency causes age-dependent impaired visual responses, and pronounced defects in rod phototransduction, with comparatively limited effects on cone function. These findings suggest that MYO1C plays a preferential role in opsin trafficking and rod photoreceptor homeostasis and may be particularly important for maintaining the functional integrity of rod OS. However, global *Myo1c*-KO mice cannot determine whether these defects arise from cell-autonomous loss of MYO1C in rods or cones. We therefore generated rod- and cone-specific *Myo1c* conditional knockout mice to further define the photoreceptor-specific functions of MYO1C. Rod-specific deletion (*Myo1c*;Rho-Cre+ mice) would determine whether loss of MYO1C in rods is sufficient to cause apo-opsin accumulation and impaired scotopic function, whereas cone-specific deletion (*Myo1c*;HRGP-Cre+ mice) would determine its role in cone opsin homeostasis and photopic function (**Supplementary Figures S4A, S4B**). These complementary models would establish whether MYO1C directly regulates opsin trafficking and photoreceptor function in a cell-autonomous manner.

### Age-Dependent Effects of Rod and Cone Photoreceptor-Specific *Myo1c* Deletion

To determine whether MYO1C is required for photoreceptor function in a cell-autonomous manner, we examined retinal function in rod- and cone-specific *Myo1c* conditional knockout mice at 2-, 6-, 8-, and 10-months of age. Rod-specific *Myo1c* deletion (*Myo1c*;Rho-Cre+ mice) resulted in a progressive decline in rod-mediated visual function with age (**Figures 8A, 8C, and 8D**). Rod function was relatively preserved in *Myo1c*;Rho-Cre+ animals at 2-months but became increasingly impaired at 6-, 8-, and 10-months, demonstrating an age-dependent requirement for MYO1C in maintaining rod photoreceptor function (**Figures 8A, 8C, and 8D**). In contrast, cone-specific *Myo1c* deletion (*Myo1c*;HRGP-Cre+) did not produce a comparable decline in cone-mediated function over the same age range (**Figures 8B, 8E–8H**). Photopic responses remained relatively preserved at 2-, 6-, 8-, and 10-months, indicating that cone photoreceptor function is substantially less dependent on MYO1C (**Figures 8B, 8E–8H**).

**Figure 8:**
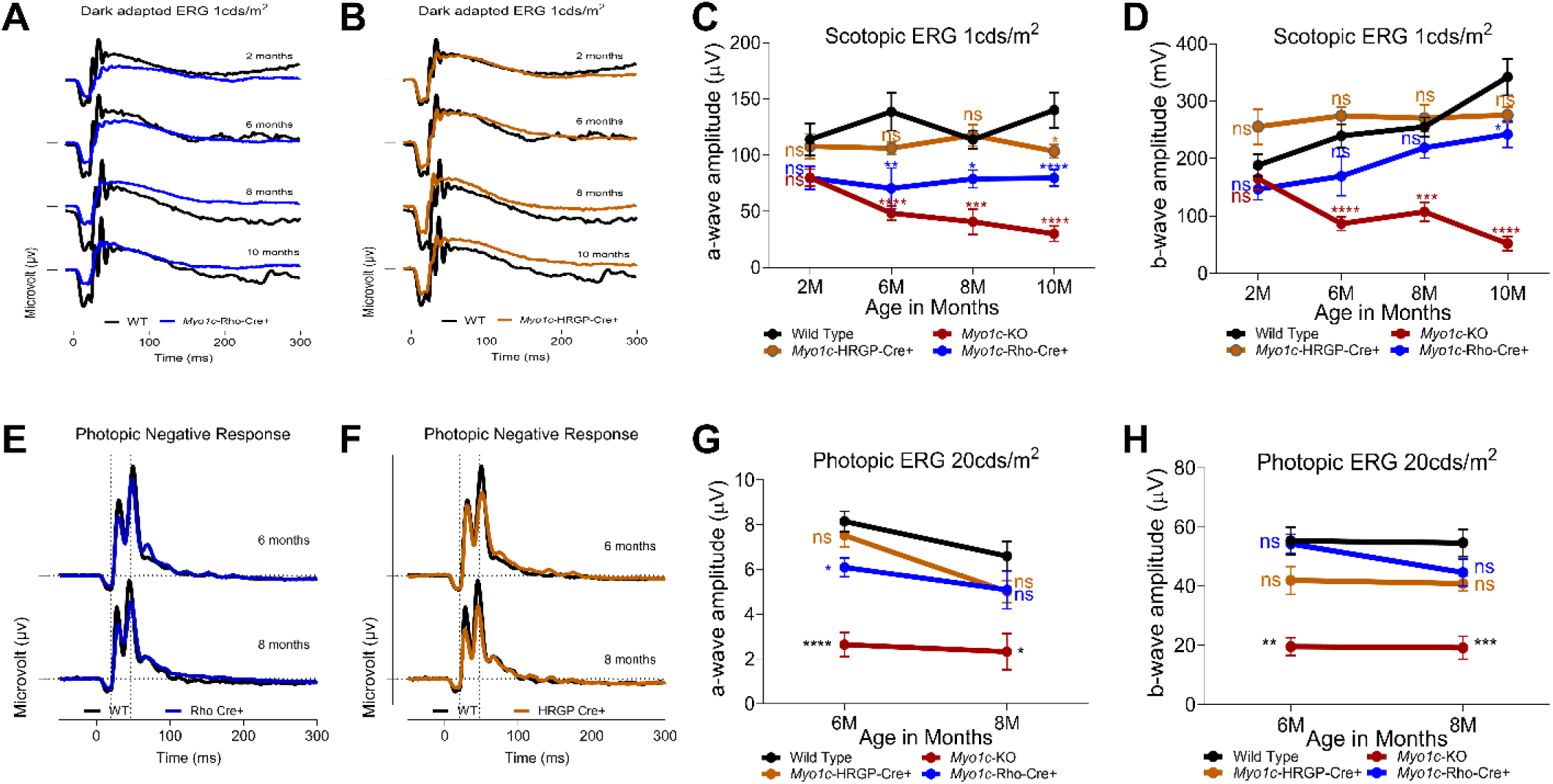
Rod-specific deletion of *Myo1c* causes age-dependent impairment of rod photoreceptor function, whereas cone-specific deletion preserves cone function. (**A-C**) Dark-adapted scotopic ERGs were recorded in response to increasing light intensities in control and rod-specific *Myo1c*-KO mice (*Myo1c*;Rho-Cre+) and cone-specific Myo1c-KO mice (*Myo1c*;HRGP-Cre+) at 2-, 6-, 8-, and 10-months of age. (**E-H**) Light-adapted photopic ERGs were recorded from control, cone-specific, and rod-specific *Myo1c*-KOat 2-, 6-, 8-, and 10-months of age. (**C, D**) Quantification of scotopic and (**G, H**) photopic ERG responses, respectively, demonstrating progressive impairment of rod-mediated function following rod-specific *Myo1c* deletion, with comparatively preserved cone-mediated function following cone-specific deletion. Data are presented as mean ± S.E.M.; statistical significance was determined using ANOVA with appropriate post-hoc comparisons. \**P* < 0.05, \*\*\**P* < 0.005, \*\*\*\**P* < 0.001.

Since photopic ERG responses showed greater variability between WT and global *Myo1c*-KO mice (**Figures 6D-6F**), we further evaluated cone function in rod- and cone-specific conditional *Myo1c*-KO mice. Photopic ERG responses were largely unchanged in both rod- and cone-specific *Myo1c* conditional knockout mice (**Figures 8E-8H**), consistent with the findings in global *Myo1c*-KO mice. These results indicate that loss of MYO1C causes progressive retinal dysfunction that is predominantly rod-mediated, whereas cone function appears to be relatively preserved.

Photoreceptor recovery following light stimulation further demonstrated this rod-selective phenotype. A-wave recovery was significantly impaired in global *Myo1c*-KO mice (**Supplementary Figure S5A, S5B)** and was recapitulated in rod-specific *Myo1c* conditional knockout mice (**Supplementary Figure S5E, S5F)**. In contrast, a-wave recovery was preserved in cone-specific *Myo1c*-KO mice (**Supplementary Figure S5C, S5D**). These findings indicate that MYO1C is required not only for normal rod photoreceptor responsiveness, but also for efficient recovery following light stimulation. The progressive decline in scotopic function following rod-specific *Myo1c* deletion, together with preserved photopic function following cone-specific *Myo1c* deletion (**Supplementary Figure S6)**, provides strong evidence for a cell-autonomous and rod-selective requirement for MYO1C in photoreceptor function and recovery.

### Preservation of Inner-Retinal Function in *Myo1c*-Deficient Mice

Because loss of MYO1C was associated with progressive reductions in ERG responses, we next sought to determine whether this functional impairment was restricted primarily to photoreceptors or also involved inner retinal circuitry. We therefore analyzed oscillatory potentials (OPs), which are generated predominantly by inner-retinal circuits and provide a functional measure of post-receptor signaling. Notably, oscillatory potentials (OPs) remained unchanged across all ages examined in global and conditional *Myo1c*-KO animals (**Supplementary Figure S7A-S7F**), indicating relative preservation of inner-retinal function despite the progressive decline in ERG responses in these mice. This finding provides a functional rationale for attributing the progressive ERG deficits predominantly to dysfunction of the outer retina, particularly photoreceptors, rather than to generalized retinal dysfunction. Collectively, these findings reveal a temporally distinct pattern of retinal dysfunction in *Myo1c*-deficient mice, characterized by progressive photoreceptor dysfunction with relative preservation of inner-retinal signaling. These functional deficits are consistent with the observed disruption of rhodopsin trafficking and rod photoreceptor organization. Together, these findings demonstrate that loss of MYO1C results in substantial impairment of photoreceptor function and visual performance.

### *Myo1c* Deficiency Selectively Disrupts Rod Photoreceptor IS-OS Integrity Without Significant Photoreceptor Cell Loss

Optical coherence tomography (OCT) and B-scan analyses of 2-month-old global *Myo1c*-KO and conditional *Myo1c*;Rho-Cre+ mice revealed reduced photoreceptor IS-OS and overall retinal thickness only in the global *Myo1c*-KO mice (**Supplementary Figures S8A-8D**). Longitudinal analysis of global and conditional *Myo1c*-KO mice at 6-months of age showed that, compared with WT mice, both global *Myo1c*-KO and rod-specific *Myo1c*;Rho-Cre+ mice exhibited a significant reduction in photoreceptor IS-OS layer thickness, with no change in photoreceptor ONL thickness (**Supplementary Figures S8A, S8E-S8G**). In contrast, no changes in photoreceptor IS-OS, ONL, or overall retinal thickness were observed in cone-specific *Myo1c*;HRGP-Cre+ mice at 6-months of age (**Supplementary Figures S8A-S8G**).

H&E analysis of retinas from 6-month-old global *Myo1c*-KO, rod-specific *Myo1c;*Rho-Cre+, and cone-specific *Myo1c;HRGP-Cre+* mice revealed a significant reduction in the photoreceptor IS-OS layer in both global *Myo1c*-KO and *Myo1c;Rho-Cre+* mice, but not in *Myo1c;*HRGP-Cre+ mice, compared with age-matched WT animals (**Figures 9A, 9B**). In contrast, ONL thickness was preserved across all three *Myo1c*-deficient groups and remained comparable to that of age-matched WT mice (**Figures 9C and 10**). These findings indicate that loss of MYO1C, particularly in rod photoreceptors, results in progressive disruption of photoreceptor architecture without significant loss of photoreceptor cell bodies at 6-months of age. These findings further suggest that *Myo1c* loss primarily compromises photoreceptor IS-OS integrity and retinal structural maintenance rather than causing substantial photoreceptor cell loss, as indicated by the preserved ONL thickness. Furthermore, the similar reduction in IS-OS thickness in global and rod-specific *Myo1c;*Rho-Cre+ mice, together with the absence of structural abnormalities in cone-specific *Myo1c;*HRGP-Cre+ mice, supports a predominant cell-autonomous role for MYO1C in maintaining rod photoreceptor structural integrity and function.

**Figure 9.**
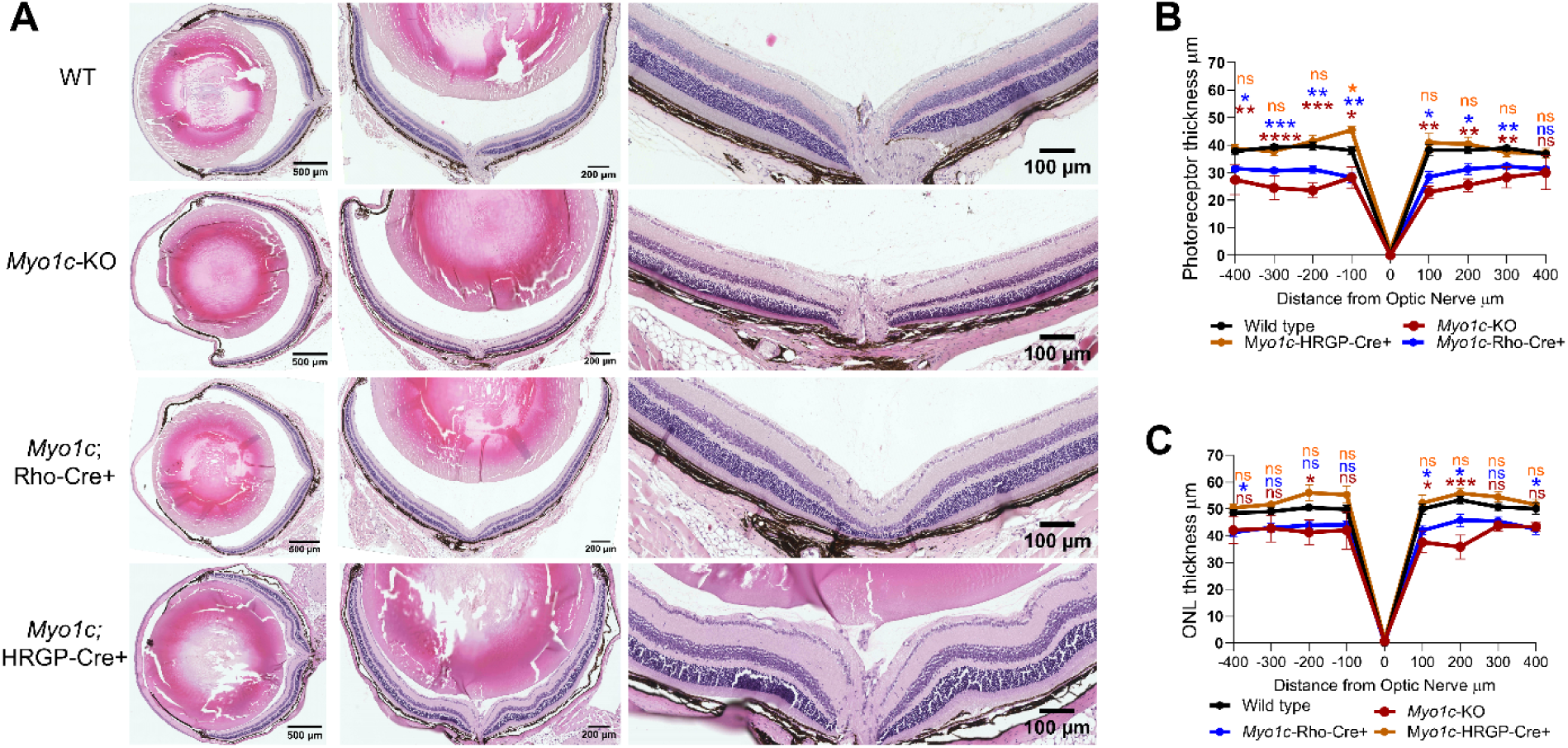
H&E staining of retinas from global and photoreceptor-specific *Myo1c* knockout mice. Representative hematoxylin and eosin (H&E) stained retinal sections from 6-month-old control/wild type, global *Myo1c* knockout (*Myo1c*-KO), and photoreceptor-specific *Myo1c* rod conditional knockout (*Myo1c*;Rho-Cre+) and *Myo1c* cone conditional knockout (*Myo1c*;HRGP-Cre+) mice. Retinal morphology and organization were evaluated across the major retinal layers, including the photoreceptor cell layer, outer nuclear layer (ONL), and total retinal thickness. No overt abnormalities in retinal architecture were observed in *Myo1c*;HRGP-Cre+knockout mice compared with controls, and global *Myo1c*-KO and Myo1c;Rho-Cre+ mice . Global *Myo1c* knockout mice exhibited [describe observed morphological changes, if applicable]. Statistical significance was determined using ANOVA. \**P* < 0.05, **P < 0.01, \*\*\**P* < 0.005, \*\*\*\**P* < 0.001; ns, not significant.

**Figure 10.**
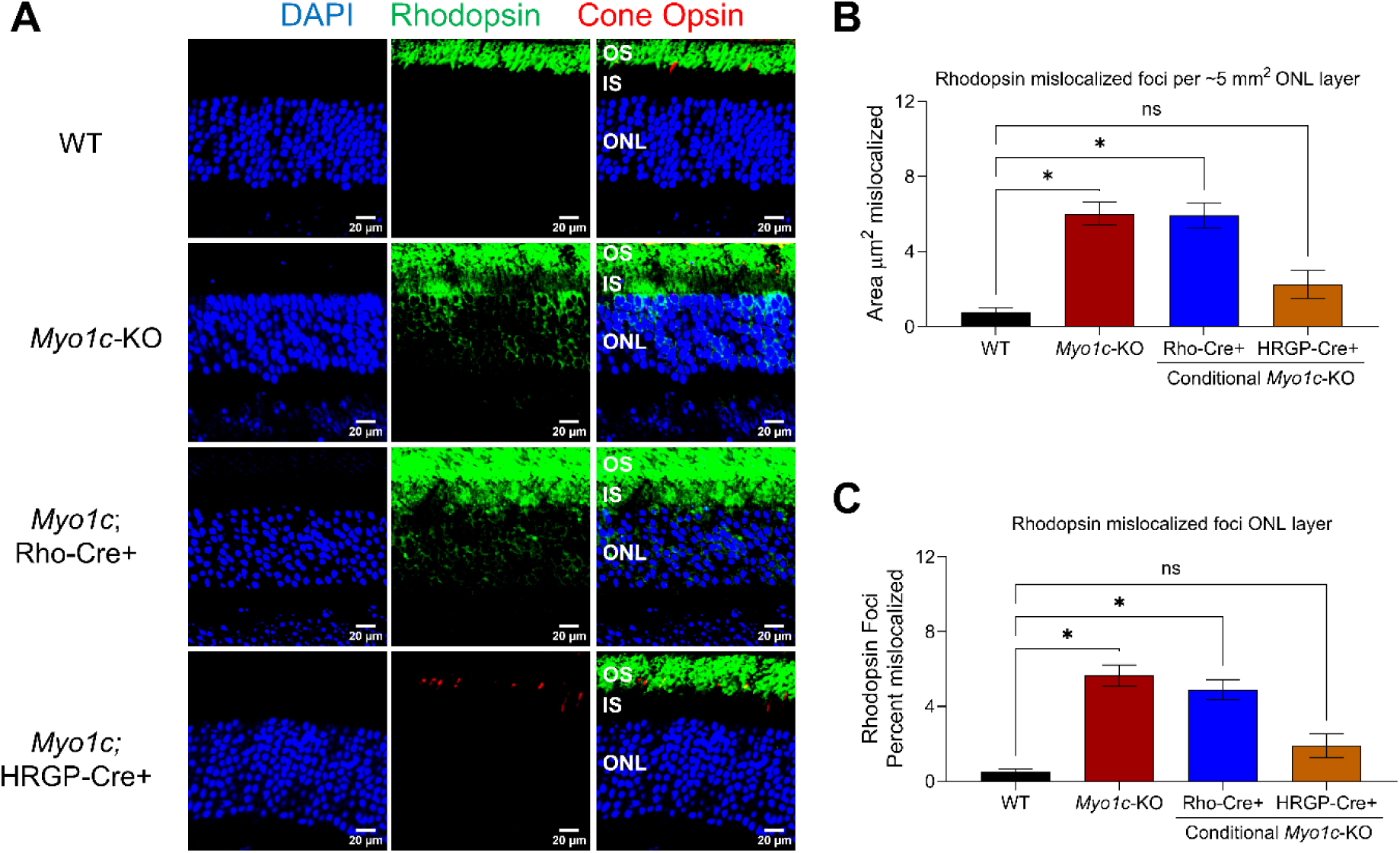
Rhodopsin and cone opsin staining in global and photoreceptor-specific *Myo1c* knockout mice. **(A)** Representative immunofluorescence images showing rhodopsin and cone opsin expression and localization in retinal sections from 6-month-old control/wild-type, global *Myo1c* knockout (*Myo1c*-KO), photoreceptor-specific rod conditional *Myo1c* knockout (*Myo1c*;Rho-Cre+), and cone conditional *Myo1c* knockout (*Myo1c*;HRGP-Cre+) mice. Rhodopsin immunostaining (1D4 antibody, green) was used to assess rod photoreceptor outer segment morphology and opsin localization, whereas cone opsin immunostaining (R/G opsins, red) was used to evaluate cone photoreceptor distribution and opsin localization. Retinal sections were examined for changes in photoreceptor organization, opsin expression, and subcellular localization. Representative images are shown for each genotype. (**B, C**) Quantification of rhodopsin fluorescence intensity and localization are shown in the corresponding graphs. Statistical significance was determined using ANOVA. \**P* < 0.05; ns, not significant.

IHC staining for rhodopsin (1D4) and red/green (R/G) cone opsins in retinal sections from 6-month-old WT (controls), global *Myo1c*-KO, rod-specific *Myo1c;Rho-Cre+*, and cone-specific *Myo1c;HRGP-Cre+* mice revealed significant rhodopsin mislocalization only in global *Myo1c*-KO and *Myo1c;Rho-Cre+* mice compared with age-matched WT animals, which was characterized by increased rhodopsin immunoreactivity within the photoreceptor cell bodies and IS, rather than its normal enrichment within the photoreceptor OS (**Figures 10A; quantified in Figures 10B, 10C**). In contrast, rhodopsin localization was largely preserved in *Myo1c;HRGP-Cre+* mice (**Figure 10A**). Similarly, R/G cone opsin localization was not significantly altered in cone-specific *Myo1c;HRGP-Cre+* mice compared with WT controls (**Figures 10A**).

Finally, fluorescein angiography analysis of global *Myo1c*-KO mice at 6 months of age showed no apparent abnormalities in the retinal vasculature compared with age-matched WT controls, indicating that loss of *Myo1c* does not cause detectable retinal vascular abnormalities at this age (**Supplementary Figure S9**). These findings demonstrate that loss of MYO1C in rods, but not cones, preferentially disrupts rhodopsin trafficking and OS organization. The concordance between rhodopsin mislocalization, reduced IS-OS thickness, and impaired ERG responses further supports a critical role for MYO1C in maintaining proper rhodopsin trafficking and rod photoreceptor function.

## DISCUSSION

Rhodopsin trafficking from the photoreceptor IS to the OS is essential for maintaining photoreceptor cell structure and function. Although several components of the rhodopsin transport machinery have been identified, the molecular motors that facilitate rhodopsin movement and membrane delivery remain incompletely understood [7, 31]. In this study, we identify using both in vitro and in vivo approaches that the unconventional myosin motor protein MYO1C acts as a previously unrecognized regulator of rhodopsin trafficking and demonstrate that its function is particularly important in rod photoreceptors. Our findings provide convergent structural, biochemical, cellular, and in vivo evidence supporting a model in which MYO1C interacts with rhodopsin through its C-terminal cargo-binding region to facilitate rhodopsin trafficking and photoreceptor homeostasis.

Our in-silico docking analysis identified a putative interaction between the C-terminal region of MYO1C and the C-terminal region of rhodopsin. Of particular interest, this region of rhodopsin contains the conserved VxPx motif, which is essential for rhodopsin trafficking from the IS to the OS. Several disease-associated rhodopsin mutations that result in abnormal intracellular localization occur within or near this region [4,8], raising the possibility that disruption of the MYO1C-rhodopsin interaction could contribute to defective rhodopsin trafficking in Retinitis Pigmentosa (RP). Although docking predictions alone cannot establish a direct interaction, the computational findings were supported by our biochemical studies demonstrating robust association between full-length MYO1C and rhodopsin. Importantly, deletion of the MYO1C C-terminal region abolished this interaction, identifying the C-terminus as a critical region for MYO1C-rhodopsin association in COS1 and hTERT-RPE1 cells.

The functional importance of the MYO1C C-terminus was further demonstrated by live-cell trafficking experiments in COS1 and hTERT-RPE1 cells. Expression of full-length MYO1C promoted efficient movement of rhodopsin-containing vesicular structures, whereas deletion of the C-terminal region resulted in prominent intracellular rhodopsin accumulation and reduced membrane-associated rhodopsin. These observations were supported by experiments in serum starved hTERT-RPE1 cells, where rhodopsin efficiently localized to the primary cilium in the presence of full-length MYO1C but showed impaired ciliary targeting following C-terminal deletion. FRAP experiments provided an additional quantitative measure of this trafficking defect. Loss of the MYO1C C-terminus substantially prolonged the recovery time of rhodopsin fluorescence within the bleached region of interest (ROI) and reduced its apparent diffusion coefficient, consistent with impaired delivery of rhodopsin from intracellular compartments to the plasma membrane. Importantly, MYO1C mobility itself was not substantially altered by C-terminal deletion, consistent with retention of the N-terminal motor and actin-binding domains. Together, these findings support a model in which the MYO1C C-terminus contributes primarily to cargo interaction and trafficking, whereas the N-terminal motor domain provides the mechanical activity via ATPase required for movement along actin-associated pathways [14,19,24,32–35].

The localization of MYO1C within isolated native murine rod photoreceptors further supports its physiological relevance in photoreceptor cell homeostasis. MYO1C was detected in both the IS and OS and was associated with rhodopsin-positive OS structures. The presence of MYO1C within these compartments is consistent with a role in the intracellular trafficking and membrane-associated organization of rhodopsin. Although our data support a role for MYO1C in rhodopsin trafficking, they do not establish whether MYO1C functions as the primary motor responsible for long-range movement of rhodopsin from the IS to the OS. Photoreceptor cargo trafficking is a highly coordinated process involving microtubule- and actin-dependent mechanisms, and MYO1C may function at a specific trafficking or membrane-delivery step rather than as the sole motor for rhodopsin transport [33–35]. Further studies will be required to define the precise stage at which MYO1C acts and how its ATPase motor activity is coordinated with other components, including F-actin [29,30], of the photoreceptor trafficking machinery.

Our in vivo findings demonstrate that disruption of MYO1C-dependent rhodopsin trafficking has important consequences for rod photoreceptor cell function. Global *Myo1c* deficiency produced an age-dependent accumulation of apo-opsin, accompanied by progressive impairment of retinal function. At 2-months of age, scotopic ERG responses were relatively preserved, whereas significant reductions in both a- and b-wave amplitudes emerged by 6-months and persisted through 10-months of age. This temporal progression suggests that loss of MYO1C does not immediately compromise photoreceptor function but instead produces a cumulative defect in photoreceptor homeostasis. The progressive accumulation of apo-opsin provides a potential mechanistic explanation for this age-dependent functional decline. Increased apo-opsin may alter the basal activity and sensitivity of the phototransduction cascade and reduce the functional reserve of rods, ultimately contributing to the progressive decline in scotopic responses [36, 37].

The functional phenotype in *Myo1c* mice was predominantly rod photoreceptor cell mediated. Global *Myo1c* deficiency produced substantial reductions in scotopic a- and b-wave responses, whereas photopic responses were comparatively preserved. This distinction became even more evident in the photoreceptor-specific *Myo1c* conditional knockout models. Rod-specific deletion of *Myo1c* recapitulated the progressive decline in scotopic function observed in global *Myo1c* knockout animals, whereas cone-specific *Myo1c* deletion did not produce a comparable decline in photopic function. These findings strongly support a cell-autonomous requirement for MYO1C in rods and indicate that rods are substantially more dependent on MYO1C function for protein transport than cones for maintenance of normal photoreceptor function. The impaired a-wave recovery observed following global and rod-specific *Myo1c* deletion provides additional evidence for a role of MYO1C in the physiological recovery of rods following light stimulation. The a-wave primarily reflects the electrical response of photoreceptors, and delayed recovery indicates that MYO1C deficiency affects not only the magnitude of the rod response but also the ability of the photoreceptor to return efficiently toward its dark-adapted state [36]. The preservation of a-wave recovery following cone-specific *Myo1c* deletion further reinforces the rod-selective nature of this phenotype. Thus, MYO1C appears to contribute to multiple aspects of rod physiology, including maintenance of rhodopsin homeostasis, phototransduction responsiveness, and recovery following activation. The selective phenotype in rod-specific *Myo1c* knockout mice, together with preserved cone function in cone-specific knockout animals, provides new evidence for cell type specific requirements for MYO1C in photoreceptor physiology. Further studies examining the molecular components and trafficking pathways underlying the delayed rod recovery will be important for defining how MYO1C supports normal phototransduction and photoreceptor homeostasis.

The preferential vulnerability of rods to MYO1C loss may reflect fundamental differences in rhodopsin abundance, rod photoreceptor OS organization, or trafficking requirements between rods and cones. Rods contain exceptionally high levels of rhodopsin, and the continuous renewal of the rod OS places substantial demands on the intracellular trafficking machinery. Even a relatively modest impairment in rhodopsin transport could therefore produce progressive accumulation of mislocalized opsin and increasing cellular stress over time. In contrast, cones may possess compensatory trafficking mechanisms or may rely less extensively on MYO1C-dependent pathways. The preservation of cone function following cone specific *Myo1c* deletion suggests that alternative mechanisms are sufficient to maintain cone opsin trafficking and phototransduction in the absence of MYO1C.

An important implication of these findings is that MYO1C dysfunction may represent a previously unrecognized mechanism contributing to retinal degeneration associated with abnormal opsin trafficking and accumulation. Mislocalization of rhodopsin is a common feature of several inherited retinal degenerative disorders, and our findings suggest that disruption of a motor protein-cargo interaction could contribute to this process. The observation that disease associated rhodopsin mutations occur within the predicted MYO1C-interacting region is particularly intriguing. It raises the possibility that some rhodopsin mutations may impair not only intrinsic properties of the rhodopsin molecule but also its interaction with trafficking machinery. Direct testing of disease-associated rhodopsin variants for MYO1C binding and trafficking efficiency will be important to determine whether this mechanism contributes to specific forms of retinitis pigmentosa.

Several limitations should be considered. First, the docking studies provide a structural prediction rather than definitive evidence of the precise binding interface between MYO1C and rhodopsin. Second, the deletion of the MYO1C C-terminus removes multiple IQ domains and therefore does not distinguish which individual region or residue mediates rhodopsin binding. Future studies using targeted mutations within the IQ1, IQ2, post-IQ, and adjacent regions will be necessary to define the minimal rhodopsin-binding domain. Third, while our live-cell and FRAP experiments demonstrate impaired rhodopsin trafficking following C-terminal deletion, they do not fully resolve the molecular steps between cargo binding, actin-dependent movement, membrane delivery, and ciliary entry. Finally, the mechanisms underlying the apparent rod-selective dependence on MYO1C remain to be determined.

In conclusion, our findings establish MYO1C as an important regulator of rhodopsin trafficking and rod photoreceptor cell function. The combination of computational modeling, biochemical interaction studies, live-cell imaging, FRAP, native photoreceptor localization, and conditional genetic deletion provides convergent evidence that MYO1C contributes to rhodopsin trafficking through its C-terminal cargo binding region. Loss of MYO1C results in age dependent apo-opsin accumulation, impaired rod phototransduction, and delayed photoreceptor recovery, while cone function is comparatively preserved. Most importantly, the recapitulation of the phenotype by rod-specific *Myo1c* deletion establishes a cell-autonomous and preferential requirement for MYO1C in rods. These findings identify MYO1C dependent rhodopsin trafficking as a previously unrecognized mechanism regulating rod photoreceptor homeostasis and provide a potential mechanistic link between defects in intracellular cargo transport and retinal degeneration.

## Abbreviations

MYO1C: Motor Protein Myosin 1C
ERG: electroretinography
KO: knockout Rho, rhodopsin
GFP: green fluorescent protein
HRGP: human red/green pigment
Cre+: Cre recombinase
ERG: electroretinography
WT: wild type
OS: outer segment
IS: inner segment
RP: retinitis pigmentosa
FRAP: Fluorescence Recovery after Photobleaching

## AUTHOR CONTRIBUTIONS

Conceptualization, G.P.L.; methodology, G.P.L., R.R.; software, R.R.; reagents, G.P.L., H.R., J.H.L., A.A.K., F.J.K; formal analysis, R.R., G.P.L., H.R., V.N.; investigation, R.R., G.P.L., S.C., R.M., V.N.; resources, G.P.L., H.R., R.M., S.C., J.H.L., F.J.K.; conditional mice breeding, G.P.L., Y.D., R.R., data curation, R.R., G.P.L.; writing original draft preparation, G.P.L.; manuscript writing, review, and editing, R.R., H.R., G.P.L., S.C, R.M., A.A.K., F.J.K., V.N.; supervision, G.P.L.; All authors have read and agreed to the published version of the manuscript.

## ACKNOWLEDGMENTS

This work was supported by NIH-NEI grant EY030889 (G.P.L.), Minnesota Lions Gift of Sight Foundation Grant (G.P.L), VitreoRetinal Foundation Grant (R.R.)., and in part by the University of Minnesota start-up funds (G.P.L.). The authors thank Dr. Deepak Nihalani (Medical University of South Carolina) for advice with *Myo1c* conditional mice breeding strategies, and Venkateshwara Dronamraju and Ahmed Sadah (University of Minnesota) for assistance with mice husbandry and genotyping.

## DATA AVAILABILITY STATEMENT

Data sharing is not applicable to this article, as no datasets were generated or analyzed in this study. Reagents and genetically modified mice lines are available from the corresponding author upon reasonable request.

## SUPPLEMENTARY FIGURES and LEGENDS

**Supplementary Figure S1.**
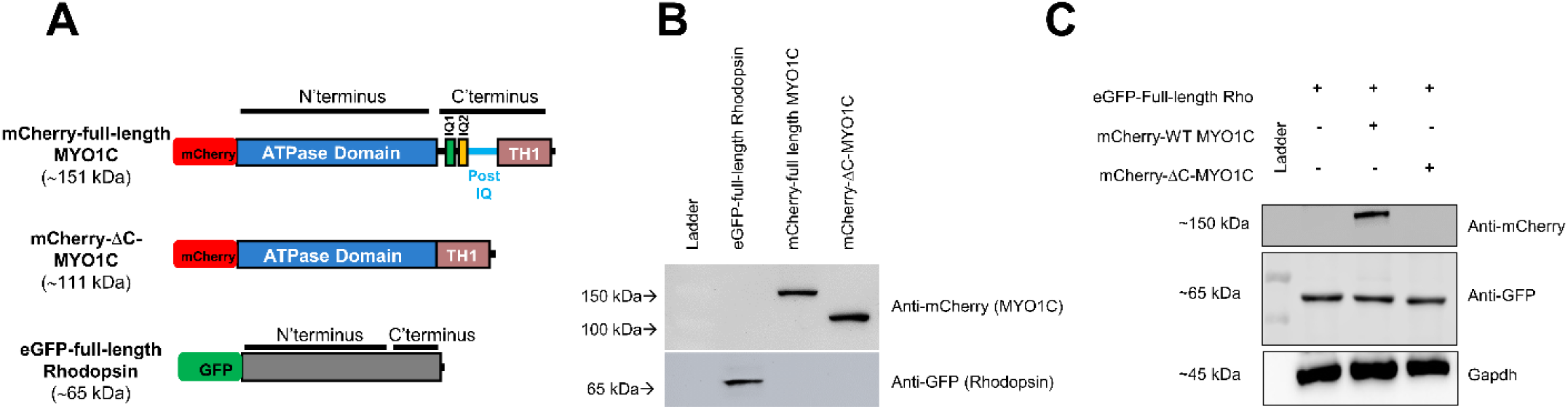
MYO1C C-terminal deletion constructs used for rhodopsin binding and trafficking studies. (**A**) Schematic representation of the mCherry-tagged MYO1C full-length and deletion constructs, along with the GFP-tagged rhodopsin construct used for binding and trafficking analyses. (**B**) Expression analysis of mCherry-MYO1C deletion constructs and GFP-rhodopsin following transient overexpression in COS-1 cells. Protein expression was confirmed by fluorescence imaging and/or immunoblot analysis prior to functional studies. (**C**) COS-1 cells were co-transfected with mCherry-tagged full-length MYO1C and GFP-rhodopsin, GFP-rhodopsin alone, or mCherry-tagged C-terminal deletion MYO1C (ΔC-MYO1C) and GFP-rhodopsin. Protein interactions were assessed by co-immunoprecipitation using an anti-GFP antibody to pull down GFP-rhodopsin, followed by Western blot analysis with an anti-mCherry antibody to detect associated MYO1C proteins. Full-length mCherry-MYO1C was detected at approximately 151 kDa, while GFP-rhodopsin was detected at approximately 65 kDa. Deletion of the MYO1C C-terminus abolished its interaction with rhodopsin, indicating that the C-terminal region of MYO1C is essential for rhodopsin binding.

**Supplementary Figure S2.**
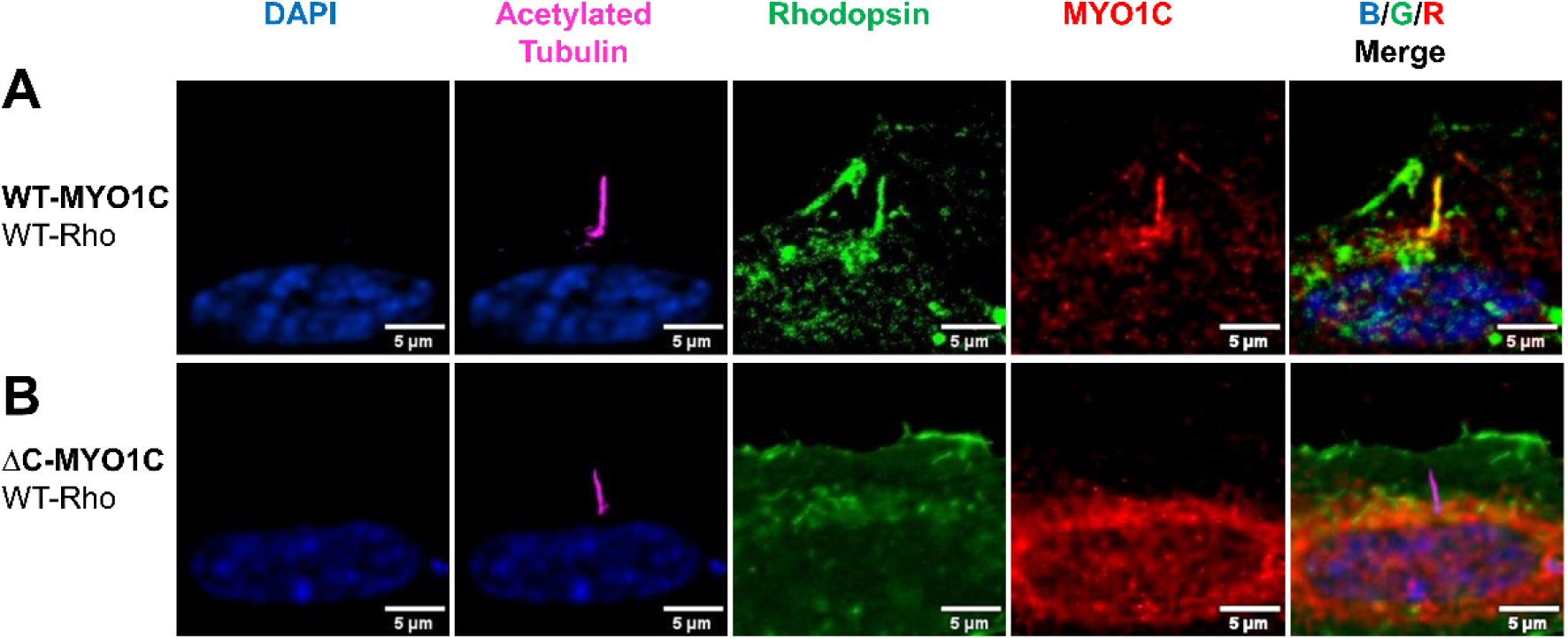
MYO1C C-terminal domain is required for Rhodopsin trafficking to the primary cilium in hTERT-RPE1 cells. hTERT-RPE1 cells were co-transfected with mCherry-WT-MYO1C or mCherry-ΔC-MYO1C (DC-MYO1C) and GFP-WT-Rhodopsin, followed by serum starvation for 24 h to induce primary ciliogenesis. (**A**) Cells expressing WT-MYO1C demonstrated trafficking of Rhodopsin-GFP to the primary cilium, with MYO1C and Rhodopsin exhibiting overlapping localization within the ciliary compartment. (**B**) In contrast, cells expressing DC-MYO1C showed impaired ciliary localization of Rhodopsin-GFP, with little or no detectable Rhodopsin within the primary cilium. Primary cilia were immunostained with an anti-acetylated tubulin antibody (magenta) to visualize the ciliary axoneme. Representative images show MYO1C (red), Rhodopsin-GFP (green), acetylated tubulin (magenta), and merged fluorescence channels. Scale bar, 5 μm.

**Supplementary Figure S3:**
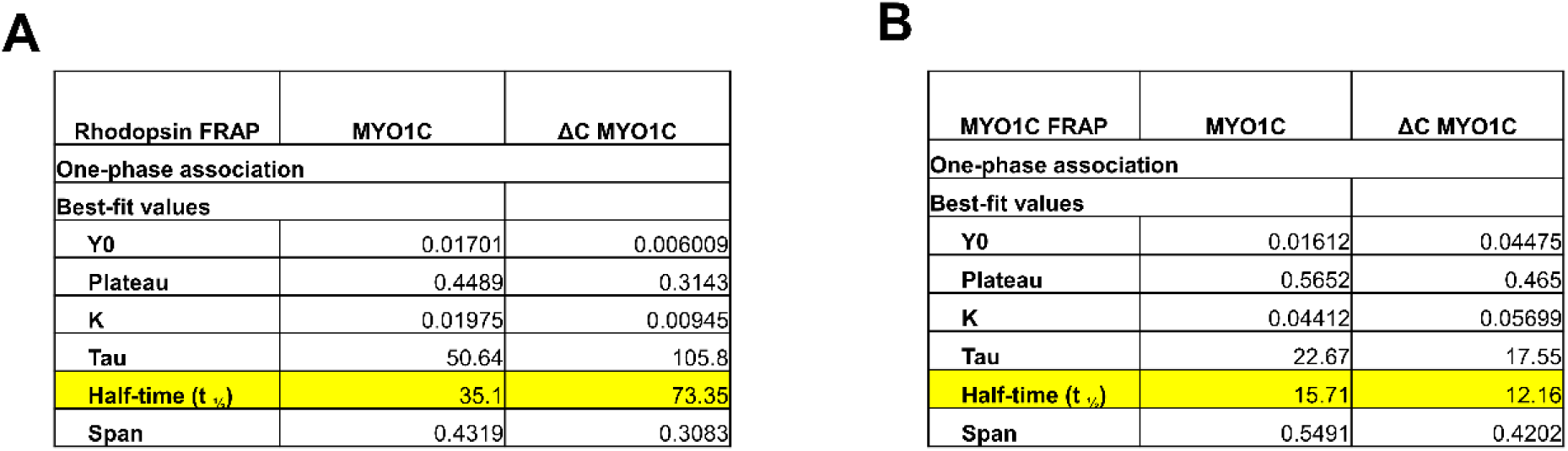
MYO1C C-terminal domain regulates rhodopsin mobility in hTERT-RPE1 cells. Fluorescence recovery after photobleaching (FRAP) analysis of (**A**) GFP-rhodopsin and (B) (B) mCherry-MYO1C in hTERT-RPE1 cells expressing WT-MYO1C or ΔC-MYO1C. The half-time of fluorescence recovery (t₁/₂), determined by exponential fitting of the recovery curves, was increased in ΔC-MYO1C-expressing cells compared with WT-MYO1C-expressing cells (73.35 vs. 35.1 s, respectively), indicating reduced rhodopsin mobility following deletion of the MYO1C C-terminal domain. Data are presented as mean values.

**Supplementary Figure S4.**
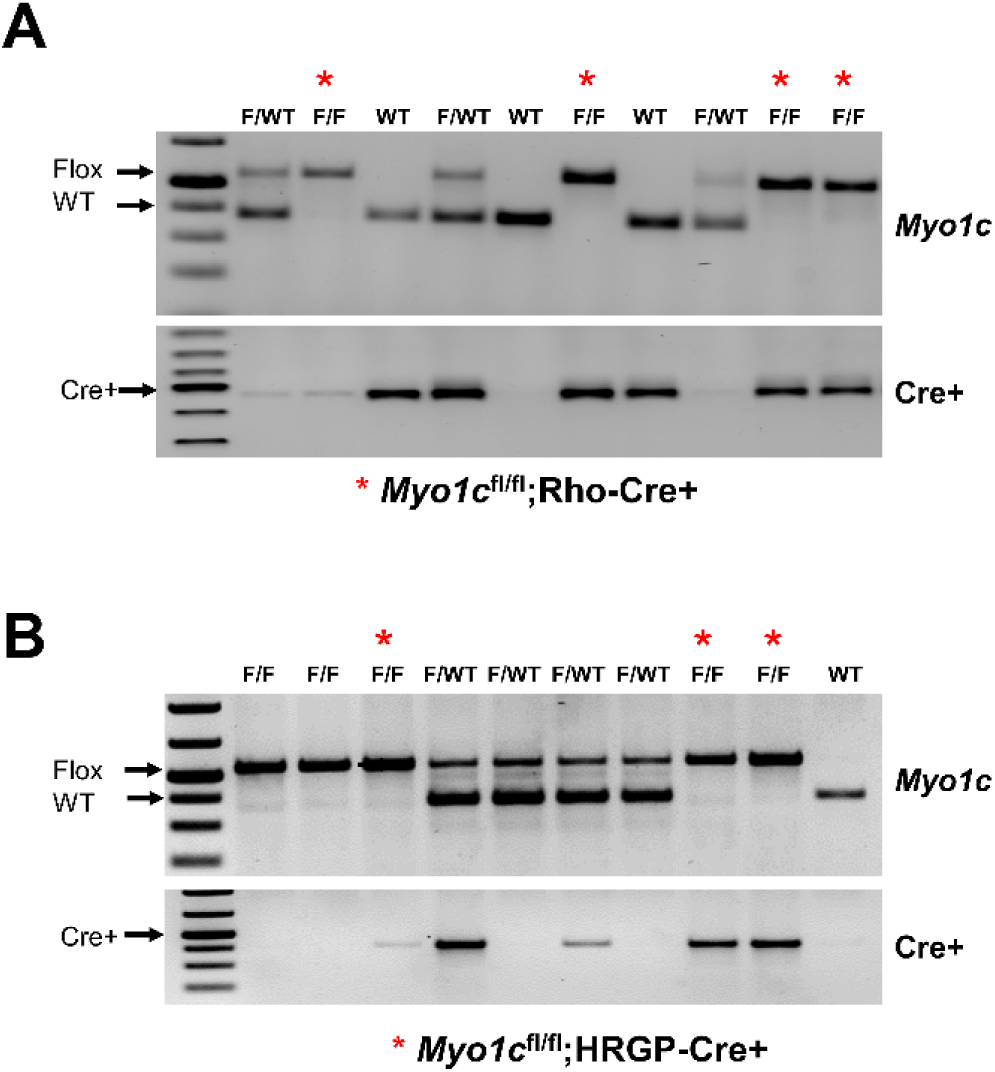
Genotyping of rod- and cone-conditional *Myo1c*-KO mice. Representative PCR genotyping results confirming the presence of the *Myo1c* floxed allele and Cre recombinase in rod- and cone-specific conditional *Myo1c* knockout mice. Genotypes were verified by PCR analysis of genomic DNA isolated from tail biopsies. The expected PCR products corresponding to the *Myo1c* floxed allele and cell-specific Cre transgenes are indicated. WT, wild type; *Myo1c*^fl/fl^, *Myo1c*^fl/wt^*, Myo1c*^fl/fl^;Cre+ recombinase. Rod-conditional *Myo1c*-KO mice were generated by crossing *Myo1c*^fl/fl^ mice with rod-specific Rho-Cre+ mice, whereas cone-conditional *Myo1c*-KO mice were generated by crossing *Myo1c*^fl/fl^ mice with cone-specific HRGP-Cre+ mice.

**Supplementary Figure S5.**
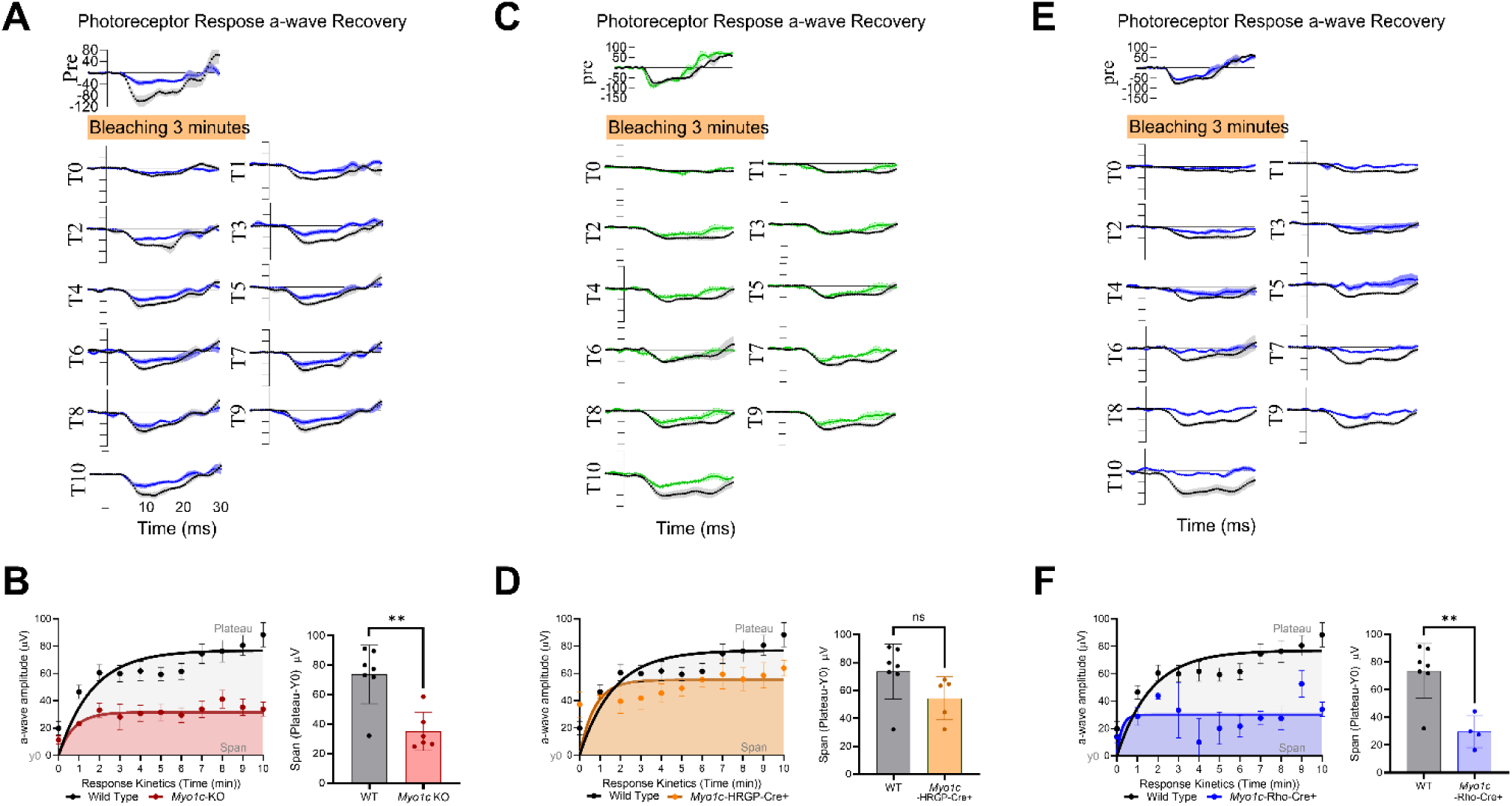
Photoreceptor response recovery in *Myo1c*-conditional knockout mice. Photoreceptor response recovery was assessed by electroretinography following exposure to a conditioning light stimulus in rod- and cone-specific *Myo1c* conditional knockout mice and corresponding control/WT mice at the indicated ages. Representative recovery traces and quantitative analysis of the recovery kinetics are shown. Global and rod-specific *Myo1c* deletion resulted in delayed recovery of photoreceptor responses (**A, C**) compared with control mice, whereas cone-specific *Myo1c* deletion had minimal effects on response recovery (**B**). Data are presented as mean ± S.E.M.; statistical significance was determined using ANOVA with appropriate post-hoc comparisons.\*\**P* < 0.01; n.s., not significant.

**Supplementary Figure S6.**
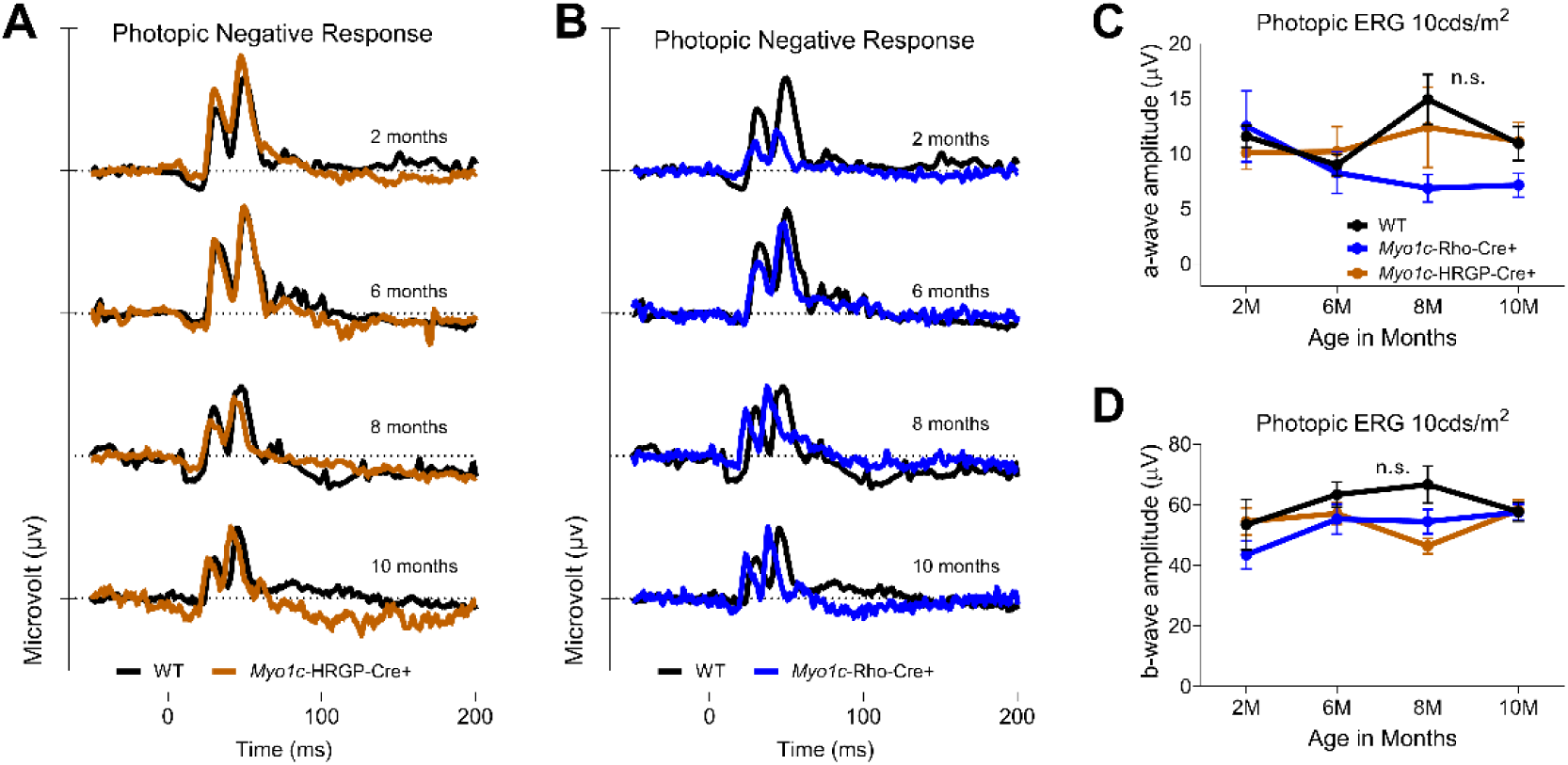
Photopic negative response ERG in *Myo1c*-KO conditional mice. Photopic negative response (PhNR) amplitudes were assessed by electroretinography to evaluate inner-retinal function in rod- and cone-specific *Myo1c* conditional knockout mice. Representative PhNR waveforms and quantitative analysis are shown for rod-specific *Myo1c* knockout (**A**), cone-specific *Myo1c* knockout (**B**), and corresponding control/WT mice at the indicated ages. PhNR amplitudes were largely preserved across the ages examined, indicating relative preservation of inner-retinal function despite the progressive decline in photoreceptor responses observed in *Myo1c*-deficient mice (**C, D**). Data are presented as mean ± S.E.M.; statistical significance was determined using ANOVA with appropriate post-hoc comparisons. n.s., not significant.

**Supplementary Figure S7.**
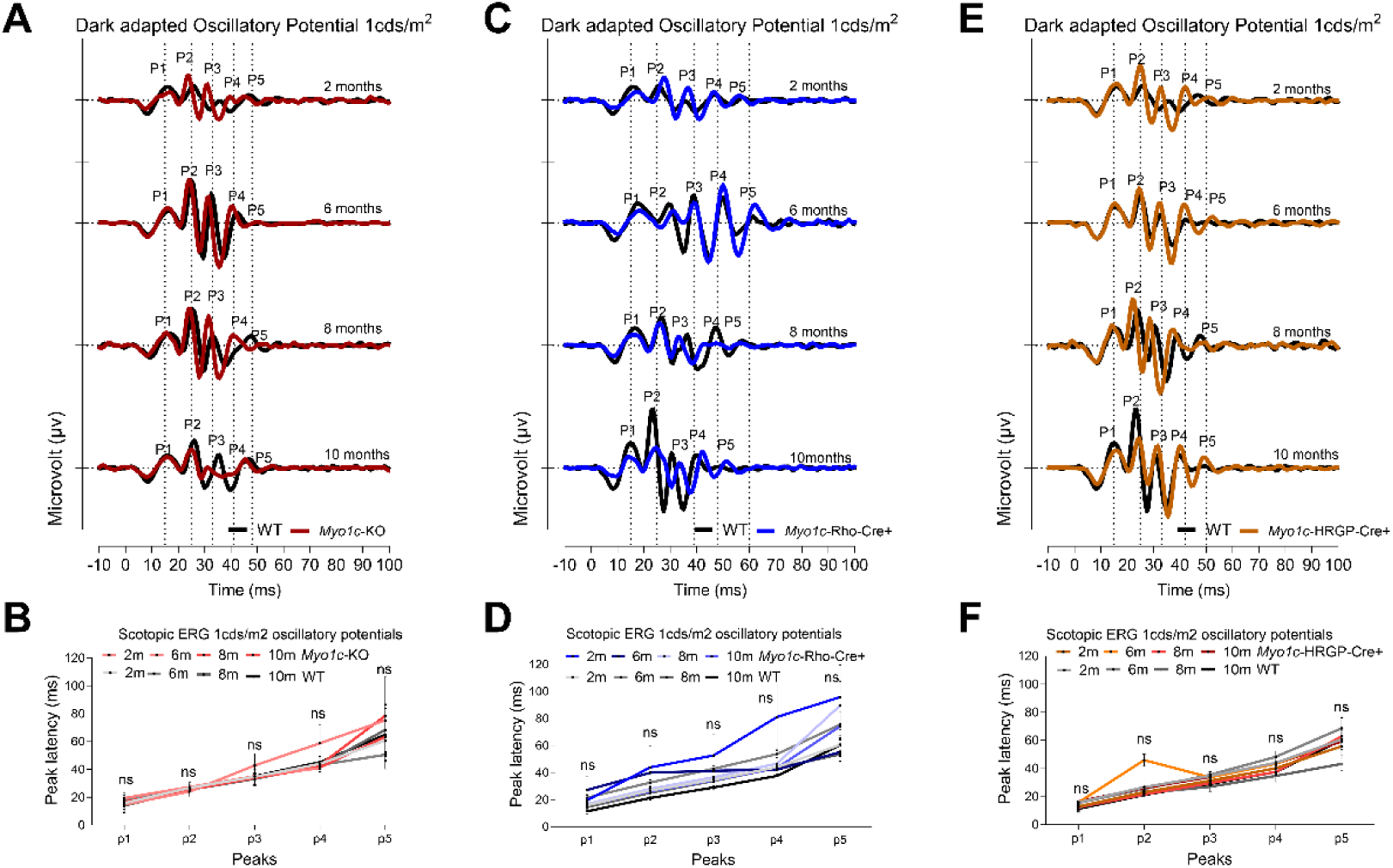
Dark-adapted oscillatory potentials in *Myo1c*-deficient mice. Dark-adapted oscillatory potentials (OPs) were recorded by electroretinography to assess inner-retinal function in global *Myo1c* knockout mice, rod-specific *Myo1c* conditional knockout mice, cone-specific *Myo1c* conditional knockout mice, and corresponding control mice at the indicated ages. Representative OP waveforms and quantitative analyses of OP amplitudes are shown. OP responses remained largely preserved across the ages examined and were comparable between *Myo1c*-deficient and control mice, indicating relative preservation of inner-retinal function despite the progressive impairment of photoreceptor responses associated with *Myo1c* deficiency. Data are presented as mean ± SEM. N.s., not significant.

**Supplementary Figure S8.**
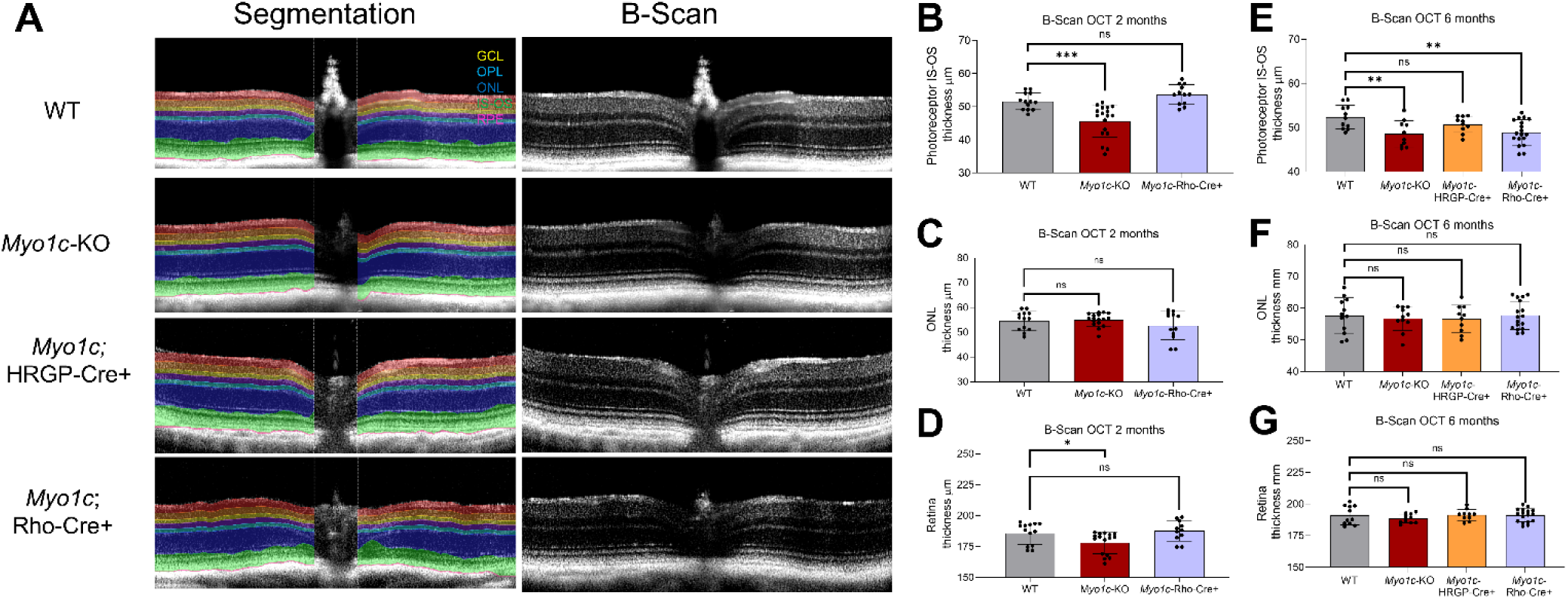
*Myo1c* deficiency selectively affects photoreceptor IS-OS integrity. Optical coherence tomography (OCT) and B-scan analyses were performed in global *Myo1c*-KO, rod-specific *Myo1c*;Rho-Cre+, cone-specific *Myo1c*;HRGP-Cre+, and WT mice. (**A**) Representative OCT images and quantification of photoreceptor IS-OS and overall retinal thickness in 6-month-old global *Myo1c*-KO and rod-specific *Myo1c*;Rho-Cre+ mice. (**B-D**) Quantification of IS-OS, outer nuclear layer (ONL), and overall retinal thickness in 2-month-old global *Myo1c*-KO and rod-specific *Myo1c*;Rho-Cre+ mice compared with age-matched WT controls. (**E-G**) Quantification of IS-OS, outer nuclear layer (ONL), and overall retinal thickness in 6-month-old global *Myo1c*-KO, rod-specific *Myo1c*;Rho-Cre+, and cone-specific *Myo1c*;HRGP-Cre+ mice compared with age-matched WT controls. Global and rod-specific *Myo1c* deletion resulted in a significant reduction in IS-OS thickness, whereas ONL thickness was unchanged. No significant differences in IS-OS, ONL, or overall retinal thickness were observed in cone-specific *Myo1c*;HRGP-Cre+ mice. Data are presented as mean ± SEM. Statistical significance was determined using ANOVA. \**P* < 0.05, **P < 0.01, \*\*\**P* < 0.001; ns, not significant.

**Supplementary Figure S9.**
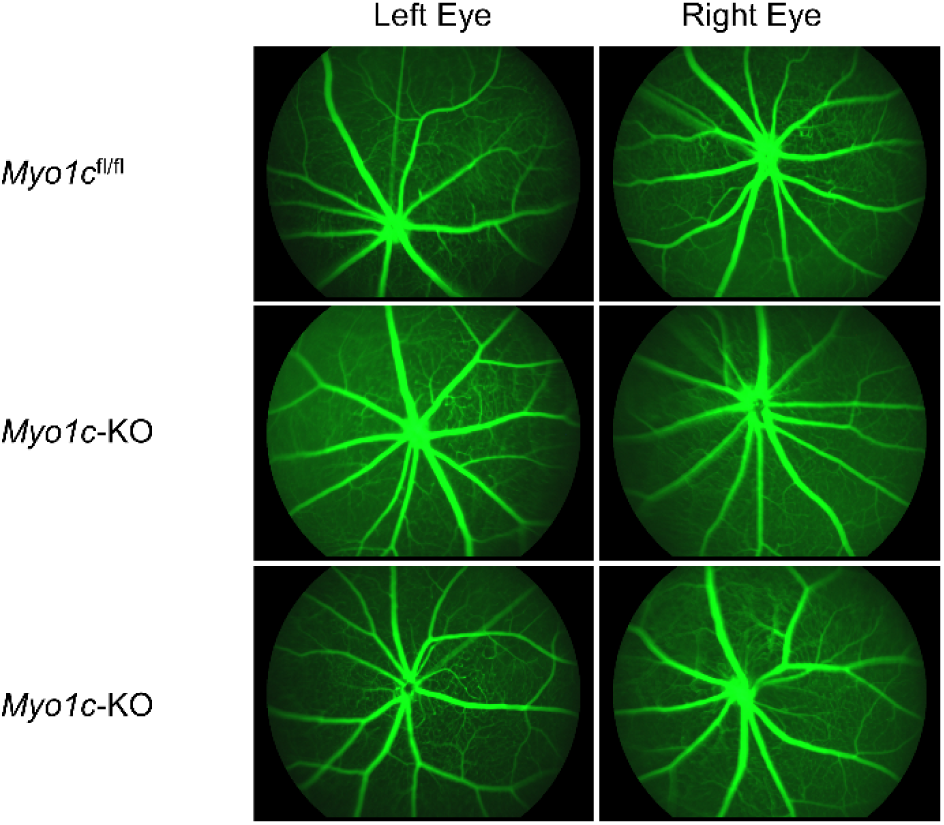
*Myo1c* deficiency does not affect retinal vascular integrity. Representative fluorescein angiography images of 6-month-old littermate control (*Myo1c^fl/fl^*) and global *Myo1c*-KO mice eyes showing comparable retinal vascular patterns, with no apparent abnormalities in vascular integrity or leakage in *Myo1c*-KO mice compared with age-matched littermate controls.

